# Selection promotes age-dependent degeneration of the mitochondrial genome

**DOI:** 10.1101/2024.09.27.615276

**Authors:** Ekaterina Korotkevich, Daniel N. Conrad, Zev J. Gartner, Patrick H. O’Farrell

## Abstract

Somatic mutations in mitochondrial genomes (mtDNA) accumulate exponentially during aging. Using single cell sequencing, we characterize the spectrum of age-accumulated mtDNA mutations in mouse and human liver and identify directional forces that accelerate the accumulation of mutations beyond the rate predicted by a neutral model. “Driver” mutations that give genomes a replicative advantage rose to high cellular abundance and carried along “passenger” mutations, some of which are deleterious. In addition, alleles that alter mtDNA-encoded proteins selectively increased in abundance overtime, strongly supporting the idea of a “destructive” selection that favors genomes lacking function. Overall, this combination of selective forces acting in hepatocytes promotes somatic accumulation of mutations in coding regions of mtDNA that are otherwise conserved in evolution. We propose that these selective processes could contribute to the population prevalence of mtDNA mutations, accelerate the course of heteroplasmic mitochondrial diseases and promote age-associated erosion of the mitochondrial genome.

## Introduction

Eukaryotic cells contain many copies of the mitochondrial genome (mtDNA). Normally all these copies are equivalent; however, largely due to replication errors or spontaneous base hydrolysis, mutations arise in individual copies creating heterogeneity or heteroplasmy^1^. To persist, a newly formed mutation must compete with the abundant copies of wild type genomes. In the female germline of *Drosophila* and *C. elegans* a PINK1-dependent intracellular quality control mechanism puts mtDNA carrying deleterious mutations under a selective disadvantage thereby limiting propagation of detrimental mutations through generations^2–5^.

While mutational accumulation in somatic tissues does not necessarily affect evolutionary stability, sustaining mtDNA quality throughout life presents challenges. An adult-human carries approximately 10^16^ mitochondrial genomes, and this huge population turns over such that about 1,000 replacement-generations occur in a lifetime. All possible simple mutations will occur billions of times and the resulting genomes will compete in an evolutionary process within our bodies^6^. Quality control could provide a purifying selection that would maintain mtDNA quality, however there is no evident purifying selection in adult *Drosophila* and cells of the human colon^7,8^. Furthermore, in multiple organisms, including mice and humans, mutant mtDNAs increase in abundance as organisms age^9–11^.

Mutations in mtDNA deleterious to oxidative phosphorylation can be masked by the co-resident wild type genomes. However, if levels of the deleterious allele rise above a critical threshold, usually in the range 60 to 90%, this protection wanes and symptoms result^12^. Thus, mechanisms influencing the cellular abundance of mtDNA mutations dictate their impact.

Levels of heteroplasmy fluctuate in cells, because mtDNA replication and segregation is random. Some simulations suggested that random chance might explain age-dependent accumulation of mtDNA mutations in somatic tissues^13^. However, any form of selection will bias outcomes and influence whether mitochondrial mutations rise to become impactful.

Work in yeast and neurospora has identified mitochondrial genomes that enjoy a selective advantage despite having negative consequences on cellular function^14,15^. These genomes acquired a replicative advantage that allowed them to out-compete the functional genomes. Work in metazoan models, *Drosophila* and *C. elegans*, similarly identified mutations with a selfish replicative advantage despite a negative effect on the host organism^16,17^. Likewise, biases in competition between diverged mammalian mtDNAs have been observed^18^. Additionally, in humans, specific mutations in the noncoding control region (NCR) climb to high abundance in particular tissues, apparently benefitting from a replicative advantage^19^. These findings suggest that a selfish ability to out replicate coresident genomes provides a general mechanism of selection. Seminal work from the Shoubridge laboratory revealed directional selection for a specific mtDNA genotype in heteroplasmic mice that favored one genotype in one tissue and the opposite genotype in another^20^. This result implicated tissue-specific nuclear genes as modifiers of the competition between two mitochondrial genomes^21,22^. This proposed nuclear influence is further supported by accumulation of specific mtDNA mutant alleles in the same tissues in different humans^19^, a genome-wide-association study in humans revealing nuclear loci associated with high abundance of particular mitochondrial alleles^23^, and direct genetic identification of nuclear modifiers of the competition between different mtDNAs in *Drosophila*^24^. Although the mechanisms by which nuclear genes alter the competition among mitochondrial genomes are unknown, nuclear genes encoding mitochondrial replication and DNA binding functions have been implicated^23,24^, and many mitochondrial alleles impacted by nuclear genes occur in noncoding sequences that are thought to have roles in mitochondrial genome replication^16,19^.

Evidence for purifying selection acting on mitochondrial genomes in the adult is limited. While little selection against a deleterious mutation was observed in adult *Drosophila*^7^, selection was observed upon introduction of additional mitochondrial stressors^7,25^. In humans heteroplasmic for a mutation in tRNA-leu(UUR), 3243A>G, the abundance of the mutant allele tends to decline in blood^26,27^ and its level is particularly low in T-cells, suggesting operation of negative selection in this population of cells^28,29^.

Although it might seem counter intuitive, evidence has been presented for selection that promotes an increase in abundance of deleterious mutations in the soma^30–34^. We refer to this proposed selection as “destructive” selection because it would raise the abundance of deleterious mutations to impactful levels, increase the population burden of mitochondrial disease alleles and enhance the severity of disease in heteroplasmic individuals.

Using single cell sequencing of liver cells from mice and humans, we have tested these ideas by analysis of the spectrum of mtDNA mutations that accumulate with age. Rather than our starting expectation that purifying selection would act to limit accumulation of deleterious mutations, we found two selective processes that accelerated the accumulation of mtDNA mutations over time. First, we identify “driver” alleles, which cause a rise in relative abundance of the affected genome within the cells in which they arise. We show that linked (‘passenger”) mutations are carried along in selective sweeps that can promote the accumulation of deleterious mutations to high levels in individual cells. Second, we found that, throughout the coding sequences of mtDNA, mutations that disrupt mitochondrial function preferentially increase in abundance, arguing for widespread action of destructive selection. We discuss how these findings help explain the bewildering variation in the progressive deterioration associated with heteroplasmic mitochondrial disease^35^.

## Results

### Numerous low abundance mutations accumulate with age in mice

In mice, *de novo* somatic mtDNA mutations seldom reach high abundance. Most are present at levels much lower than 0.1% of the total mtDNA in a tissue, which is below detection capabilities of regular Illumina sequencing. We reasoned that each specific mtDNA mutation would occur in only a tiny fraction of the cells, but in these cells and their descendants, the relative abundance of the mutation would be much higher (∼1,000-fold) than in the whole tissue and thus readily detectable with standard next-generation sequencing methods (Figure 1A). To extend sensitivity of allele detection, we developed high-throughput sequencing methods to profile mtDNA mutations in single cells (Figures 1B, S1 and S2).

**Figure 1.**
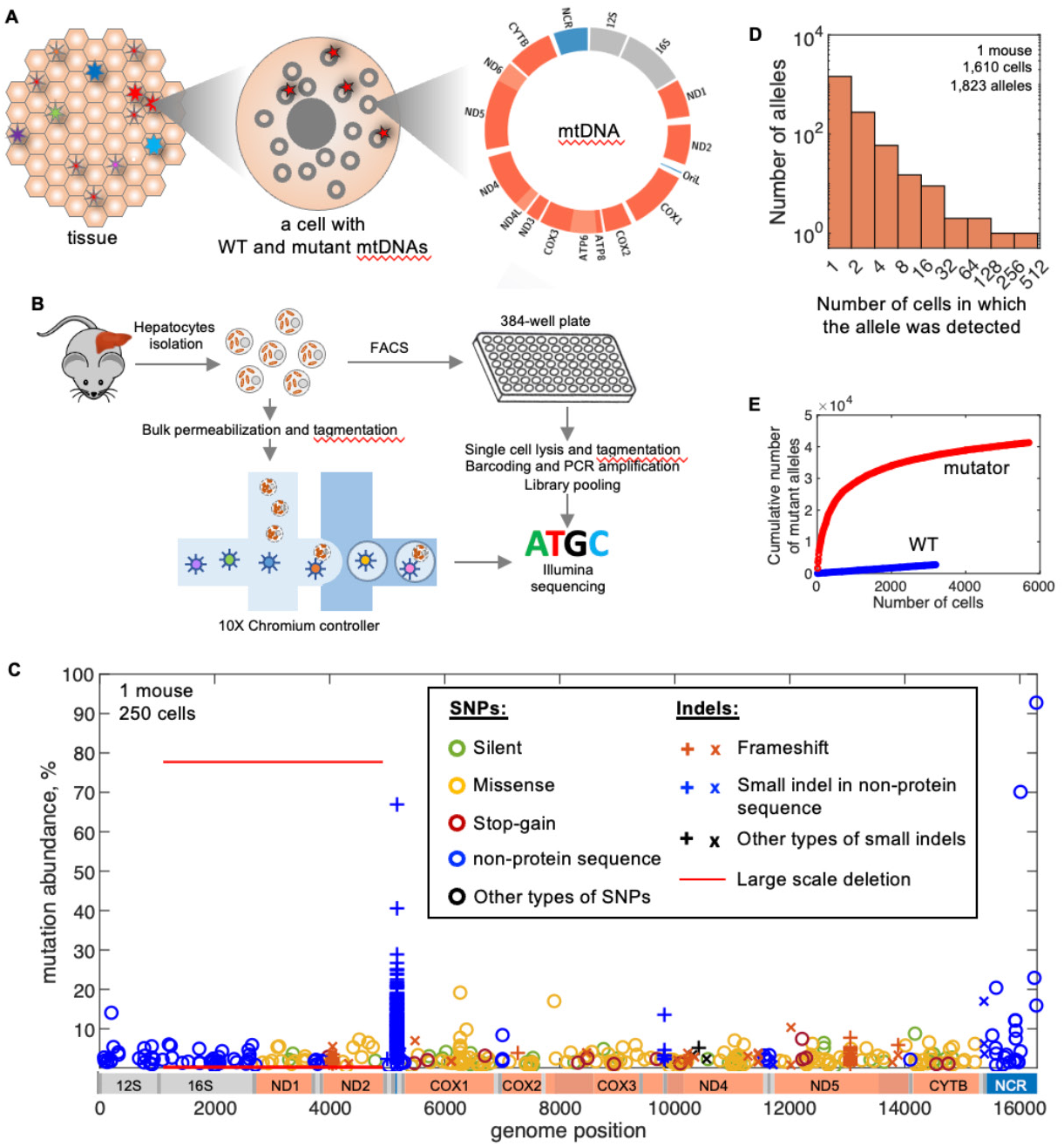
Single cell sequencing for profiling *de novo* somatic mtDNA mutations. (A) *De novo* somatic mtDNA mutations occur infrequently so that each allele is generally present in a few cells of a tissue. (B) Schematic of steps in plate-based and 10X-based single cell mtDNA sequencing to profile mtDNA mutations. (C) Spectrum of mtDNA mutations in 24-month-old C57BL6/J mouse liver. Distinct symbols indicate allele type, each occurrence is represented by a symbol indicating genomic position (X-axis) and abundance/percent of reads (Y-axis)(see Figure S1D for control) in the cell in which the mutation was recorded. Data from 250 cells are aggregated in the plot. (D) Frequency of mutant alleles detection in the cells of 24-months-old C57BL6/J mouse liver. Most observed alleles are seen in one or few cells, but rare alleles are found in most or even all the cells. (E) The number of distinct mutant alleles identified increases with number of cells analyzed. An analysis of 5,701cells from three 24-month-old heterozygous mutator mice detected 1,209,103 mutations representing 41,273 distinct alleles. Analysis of 3,195 cells from three similarly aged C57BL6/J WT mice detected 14,581 mutations representing 2,746 alleles. Data in C were generated with plate-based approach, data in D and E were generated with 10X-based approach.

To assay mtDNA sequences, we employed ATAC-seq^36^. Although this technique is commonly used to assess chromatin accessibility, it also provides a simple workflow to generate libraries enriched in mtDNA sequences^37,38^ (Figures S1B and S2C). We coupled this with cell sorting, or the 10X Genomics platform, to analyze thousands of single cells (Figures 1D and 1E), and we increased our ability to profile multiple samples at once by using sample-specific barcodes adapted from a strategy previously introduced to allow multiplexing of samples in single-cell sequencing^39^ (Figure S2; Conrad et al., in preparation). Figure 1C shows the results from 250 hepatocytes representing a larger data set from a 24-month-old C57BL6/J mouse. Most alleles are detected in less than one cell out of 1,000 (Figure 1D), and most alleles are present at very low abundance in the cell in which they are detected (Figure 1C and S3).

To increase the depth of analysis, we profiled mtDNA mutations in heterozygous mutator mice that have a proofreading-deficient mtDNA polymerase^40,41^ (Figures S3C and S3D). These mice exhibit error-prone replication, but lack the recessive premature aging phenotypes seen in the homozygote. The mutator allele was introduced from the male to avoid a contribution of maternal mtDNA mutations. Compared to WT, the heterozygous mutator mice accumulated more distinct mtDNA mutations, nearly saturating mtDNA with distinct alleles (Figure 1E). As expected for a proof-reading defect, the increase in single nucleotide polymorphisms (SNPs) was especially high (169-fold), compared to insertions (4-fold) or deletions (15-fold) (young mice, Figure S3E). Like WT mice, most mutations in the heterozygous mutator mouse exhibited a low abundance in individual cells (Figure S3). Thus, once emerged, mutations behaved much as they do in WT mouse, and persisting wild type alleles ought to promote tolerance of the numerous low abundance mutations.

To facilitate interpretation of mutational spectra, we simulated accumulation of mtDNA mutations using population genetics software (SLiM^42^) and parameters informed by our data. This allowed us to gauge how variables such as time, mutation rate, selection and mtDNA copy number influence the cellular distribution of mutations (Figure 2).

**Figure 2.**
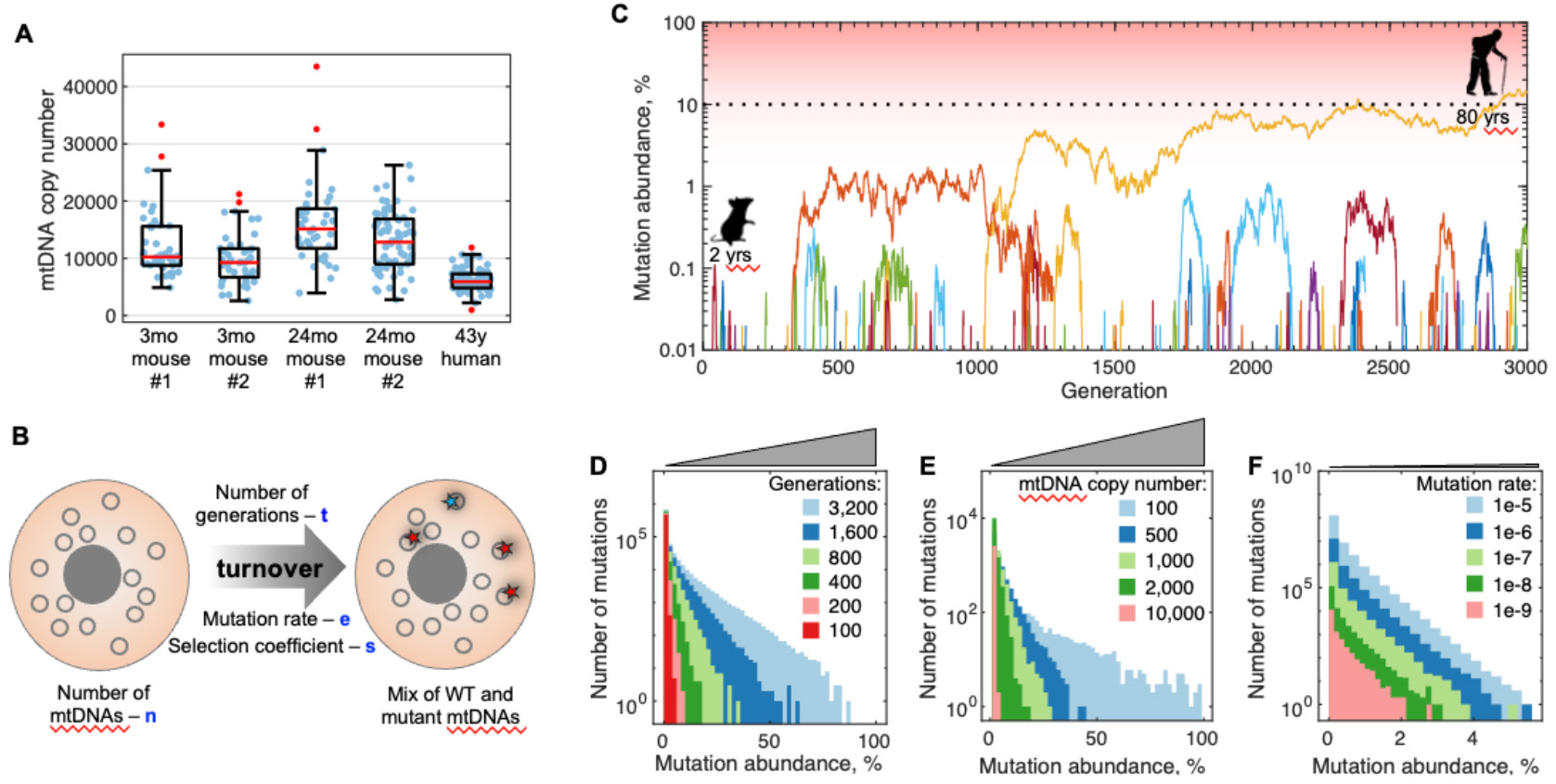
Neutral *de novo* mtDNA mutations fluctuate in abundance but high abundance mutations are infrequent, and their likelihood increases slowly with age. (A) Droplet digital PCR (ddPCR) measurements of mtDNA copy number (blue points) in single hepatocytes from young and old mice and a middle-aged human. Boxes indicate the 25th and 75th percentiles, red line marks the median. The whiskers extend to the most extreme data points not considered outliers (conventionally defined as outside 1.5 times the interquartile range above the upper quartile and bellow the lower quartile; red points). (B) Simulation of mtDNA mutations accumulation. MtDNAs were treated as individuals with a measured population size (n) in each cell, with other variables (blue) assigned. See methods section for full description of the parameters. (C) Dynamics of accumulation of simulated neutral *de novo* somatic mutations. The plot tracks the fate of a generic allele as mutants emerge in many simulations. 234 mutations (colored lines) emerged in 250 simulations. Most disappeared shortly after emergence. Only 2 persisted at the end and only one reached an abundance of 10% (black dotted line). Model parameters: mutation rate 3.16×10^-8^ per base pair per replication and 10,000 genomes per cell. (D, E, F) Simulations illustrating the impact of variables on the abundance distribution of mutations: time (number of generations) (D), mtDNA copy number (E) and mutation rate (F). Grey wedges highlight the difference in X-axis scale for D-F panels. In the lifetime of a mouse (∼80 replacement generations of mtDNA) chance accumulation of a mutation to critically high levels (usually 60%) in a cell with high mtDNA copy number is exceedingly unlikely. Models’ parameters unless specified otherwise in the figure panel: 16,299bp genome, 10,000 genomes per cell, 3.16×10^-8^ mutation rate, 80 generations, 10,000 simulated cells.

When a mutation first emerges, it is present as a single copy per cell amid about 10,000 copies of mtDNA in a mouse liver cell (Figure 2A) and is not detectable by our current methods. In a neutral model, the abundance of newly emerged mutations will fluctuate over time with the only stable outcomes being loss or fixation. Given that it starts as one out of 10,000 mtDNAs in mouse hepatocytes, the likelihood of fixation of the mutant allele is 1/10,000, with loss being by far the predominant fate^43^. Furthermore, simulations show that the rare mutations that rise to high abundance do so only after many cycles of turnover (Figures 2B-2D). In an organism with a short lifespan, such as a mouse, somatic mutations emerging in a cell with high mtDNA copy number (e.g., ∼10,000 in mouse hepatocytes) at a rate of 10^-9^ – 10^-5^ per base pair per replicative cycle and neutral behavior are exceedingly unlikely (probability <10^-5^) to climb above 10% (Figures 2E and 2F). Because mutations that impact function must climb to high abundance, we are especially interested in understanding what might promote the rise of some alleles.

We compared the spectrum of mtDNA mutations in livers of aged (24 months) and young (3 months) C57BL6/J mice (Figures 1C, S3A and S3B). The number of detected mutant mtDNA alleles increased from 3 months to 24 months of age (Figure S3). Most alleles were present at low abundance at both ages, but not all. Alleles detected in many cells, but only in a single mouse, represent clonally expanded mutations that emerged early in development (e.g. one missense allele in Figure S3A). Alleles detected in many cells of all mice represent mutations at sites that are highly mutable. These tend to be small indels in a homo-polymeric sequence (e.g. in the L origin of replication p.5172-5182). Additionally, large-scale deletions which have complex behaviors sometimes rising to high levels (Figures 1C and S3A-S3D). Beyond these special cases, many SNPs and simple indels also increase in level. Our analysis of this latter group describes a directional rise in abundance that provides evidence for two types of selective pressures, one that acts powerfully on very select alleles and one that has a weaker but widespread action.

### Mutant alleles differ in their behaviors

Sites in mtDNA are expected to differ in mutation frequency and the mutant alleles are likely to have different impacts on selection. These allele specific features impact their “behavior” in single cell data. High mutability has its predominant impact on the number of cells in which the mutation occurs while positive selection has its predominant impact on the abundance of the mutation in the cells in which it occurs. We use two approaches to relate allele behavior to selection and to mutation rate.

Graphing the average cellular abundance of an allele (AAA) in cells where it is detected versus the number of cells (C#) in which it is found produces an AAA vs C# scatter plot (Figure 3A) that gives an overview of allele-behavior. Alleles with a high mutation rate occur in more cells, with increasing mutation rates approximating a rising curve to the right on a AAA vs C# plot (simulation in Figure 3B, and grey line in Figure 3A). In contrast, alleles that are primarily influenced by positive selection rise to a high level of cellular abundance and cluster toward the top of the AAA vs C# plot (e.g. simulation in Figure 3C, and NCR alleles in Figure 3A). While statistical variance results in shifts in positions of a given allele, comparison of AAA vs C# plots from different mice show that alleles clustering in one locale on one AAA vs C# plot tend to cluster in a similar locale in an independent AAA vs C# plot (Figures 3D-3G) indicating that position on an AAA vs C# plot reflects allele specific behavior.

**Figure 3.**
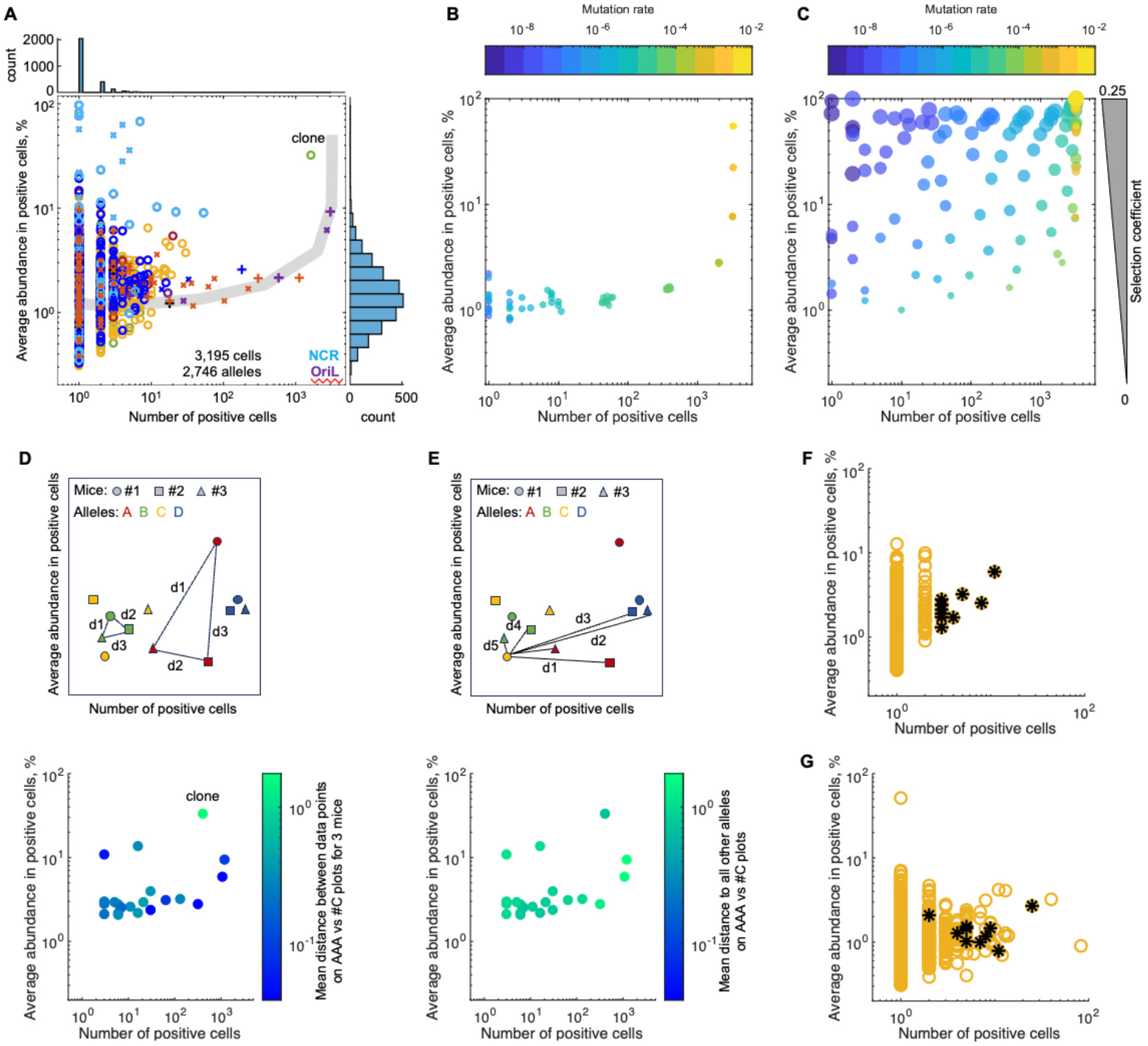
Allele behavior on AAA vs C# plots. (A) An AAA vs C# scatter plot showing the average cellular abundance of each mutant allele (AAA) in the positive cells (Y-axis), versus the number of cells (C#) in which the allele was detected (X-axis) with histograms: abundance distribution of data points (right) and distribution of data points versus cell number (top). Data shown for 3,195 cells from three 24-month-old C57BL6/J mouse livers. Symbols are as in Figure 1C except that alleles in the NCR and OriL are colored with cyan and purple, respectively. Grey line shows expected location of neutral mutations emerging with varying rates. (B, C) Simulations showing positions of neutral alleles emerging with differing mutation rates (B) or alleles differing in both mutation rate, and selection coefficient (C) on AAA vs C# plots. Mutation rate and selection coefficient indicated by color and size scales, respectively. Simulation parameters: 10,000 genomes/cell, 80 generations, 3,195 cells. (D, E) Position on an AAA vs C# plot is an allele specific property. (D) Schematic (top panel) shows four alleles (colored) that were detected in three matched mice (symbols). Distances between the three data points on the AAA vs C# plots for each mouse were measured to obtain an average mean separation (MS) as a measure of the correlation in the positions in independent mice. The bottom panel shows an AAA vs C# plot for alleles detected in all three mice and the alleles are colored according to the measured MS. (E) Unrelated alleles show a high mean separation. For each allele we measured the distance to all other unrelated alleles (schematic, top panel) and plotted the same alleles as shown in D colored according to the unrelated mean separation (bottom panel). For this analysis data from each cell were subsampled to 100,000 reads mapping to mtDNA and equal number of cells from each mouse was analyzed (n = 400 cells). (F, G) A group of non-synonymous (NS) alleles in one locale in an AAA vs C# plot from one 24-month-old C57BL6/J mouse (F) shows biased localization in AAA vs C# plot of another 24-month-old mouse from an independent experiment (G). Data presented in this figure were generated with 10X-based approach.

If, instead of just using average allele abundance, we record the abundance of a specific allele in each cell in which it is detected, we get an abundance distribution providing more detailed information. Simulations of alleles having no selection (neutral), but different mutation frequencies give a spectrum of distributions distinct from those produced when different selective forces are assigned to alleles (Figure S4A). Making the simplifying assumption that mutation rates and selective forces are constants for each allele, we used simulations to test parameter combinations to identify, at least approximately, the mutation rates and strength of selection for different alleles (Figures S4, 4C and 4D).

### Mutations conferring a selfish advantage

Certain NCR alleles were detected repeatedly, typically at high cellular abundance (Figures 3, 4A, 4C and S5). They were frequent in aged but rare in young mice (Figures 4A, 4B, S3 and S5). A search of parameter space showed that adjustments of mutation rate alone could not recapitulate the cellular distributions of this group of NCR mutations, while inclusion of a competitive advantage for the mutant genome resulted in distributions that closely resemble the data (Figures 4D and S4B). These findings suggest that many of the recurrent and especially abundant NCR mutations conferred a competitive advantage to the mitochondrial genomes on which they emerged.

**Figure 4.**
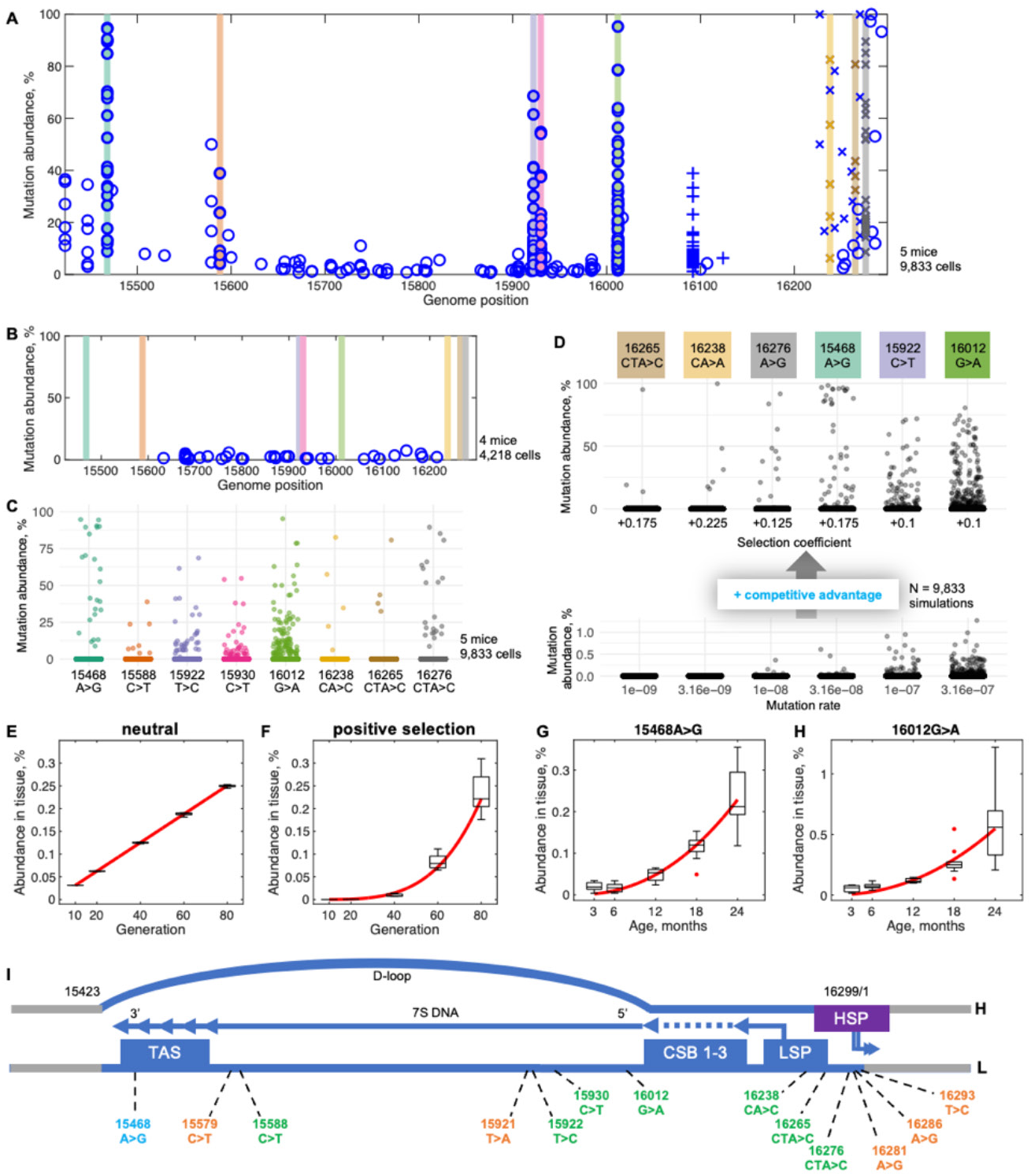
Specific mutations in NCR confer a competitive advantage. (A) The NCR region of 24-month-old WT mice is characterized by mutations that reach exceptionally high cellular abundance. Alleles present above 20% in at least 1 cell in at least 3 out 5 mice are highlighted with differently colored vertical bars. (B) The exceptional alleles (colored bars as in A) were not detected in four 3-month-old WT mice suggesting that they come to predominate with age. (C) Abundance distribution of highlighted NCR mutations among liver cells of 24-month-old WT mice. (D) Simulations show that varied mutation frequencies (lower panel) fail to mimic the abundance distributions shown in (C), while inclusion of positive selective coefficients (specified in the figure, upper panel) yields distributions resembling the data. Note the difference in Y-axis scale. (E) Simulations (n=5) ran for different numbers of cycles (proportional to age) show linear tissue level accumulation of a neutral allele. Mutation rate = 3.16×10^-5^/base pair/cycle, and mtDNA copy number = 10,000. (F) Simulations (n=5) show that a positively selected (coefficient = +0.175) mutant allele accumulates at the whole tissue level at an accelerating rate. Mutation rate = 3.16×10^-8^/ base pair/generation, and mtDNA copy number = 10,000. (G, H) Accumulation dynamics of the indicated driver alleles as measured by allele-specific ddPCR assays in bulk liver of WT mice. N = 5 mice per time point for each allele tested. (E-H) Box plots show simulated or ddPCR data. Red lines show linear (E) and power (F-H) function fitting, R^2^= 0.999, 0.945, 0.834 and 0.617, respectively. (I) Driver mutations are localized in the NCR (blue) in association with sequences thought to govern mtDNA replication: the termination associated sequence (TAS); the conserved sequence boxes (CSB1-3); the light strand promoter (LSP) and the heavy strand promoter (HSP). The LSP initiates an RNA (arrow) that primes DNA synthesis within the CSBs. DNA synthesis continues to a pause point in TAS and can be continued to promote a synthesis of new heavy strand. The allele labeled in cyan showed selective amplification in both WT and heterozygous mutator mice, whereas, at least using stringent criteria to identify drivers, the alleles labeled in green were only seen as a driver in the WT, and alleles labeled in orange were only seen as drivers in the heterozygous mutator line. Data in A-D were generated with 10X-based approach.

Simulations of mutation levels in bulk tissue show that accumulation of neutral alleles is linear, while accumulation of positively selected alleles follows a power function (Figures 4E and 4F). We developed ddPCR assays to measure abundance of 15468A>G and 16012G>A mutations in bulk mouse liver and found that tissue-level abundance of these alleles increased with age in WT mouse liver closely following a power function (Figures 4G and 4H). These observations further suggest that the recurrent mutations in the NCR are positively selected. The positively selected alleles are clustered in the NCR region in mouse and associated with DNA sequences contributing to replication (Figure 4I). As was previously argued from such associations in human^19^, we suggest that the positive selection results from a replicative advantage incurred by the mutant genomes.

### Passenger mutations

Individual cells carrying positively selected NCR-mutations occasionally had other mutations at similarly high levels (Figure 5). We suggest that in such cases a positively selected NCR allele emerged on a genome with an existing mutation. The NCR allele then acted as a “driver” promoting the relative abundance of itself and of the linked “passenger” allele. In two of the three cells shown in Figure 5, the NCR-mutation and its passenger were present at >50% of total mtDNAs, and therefore must co-reside on at least some of the mtDNAs (Figures 5A and 5C). In the third example, linkage of one of several candidate passengers is documented by individual sequencing reads recording both the NCR mutation and the nearby passenger (Figure 5G). Since associations between a driver and particular passenger are exceedingly rare, examining the abundance of “driver” and “passenger” alleles in other sequenced cells should show their independent behavior (Figures 5B, 5D and 5F). Driver alleles were found at abundant levels in multiple cells indicating an autonomous advantage. In contrast, when the “passenger” allele occurred on its own, it was found at low abundance. This is consistent with a conclusion that in cells presented in Figure 5 the “passenger” alleles gained an advantage by a chance association with a driver allele. Notably, associations of apparently deleterious mutations with a driver mutation led to their rise in abundance to levels that could impair cellular metabolism (Figures 5A and 5C).

**Figure 5.**
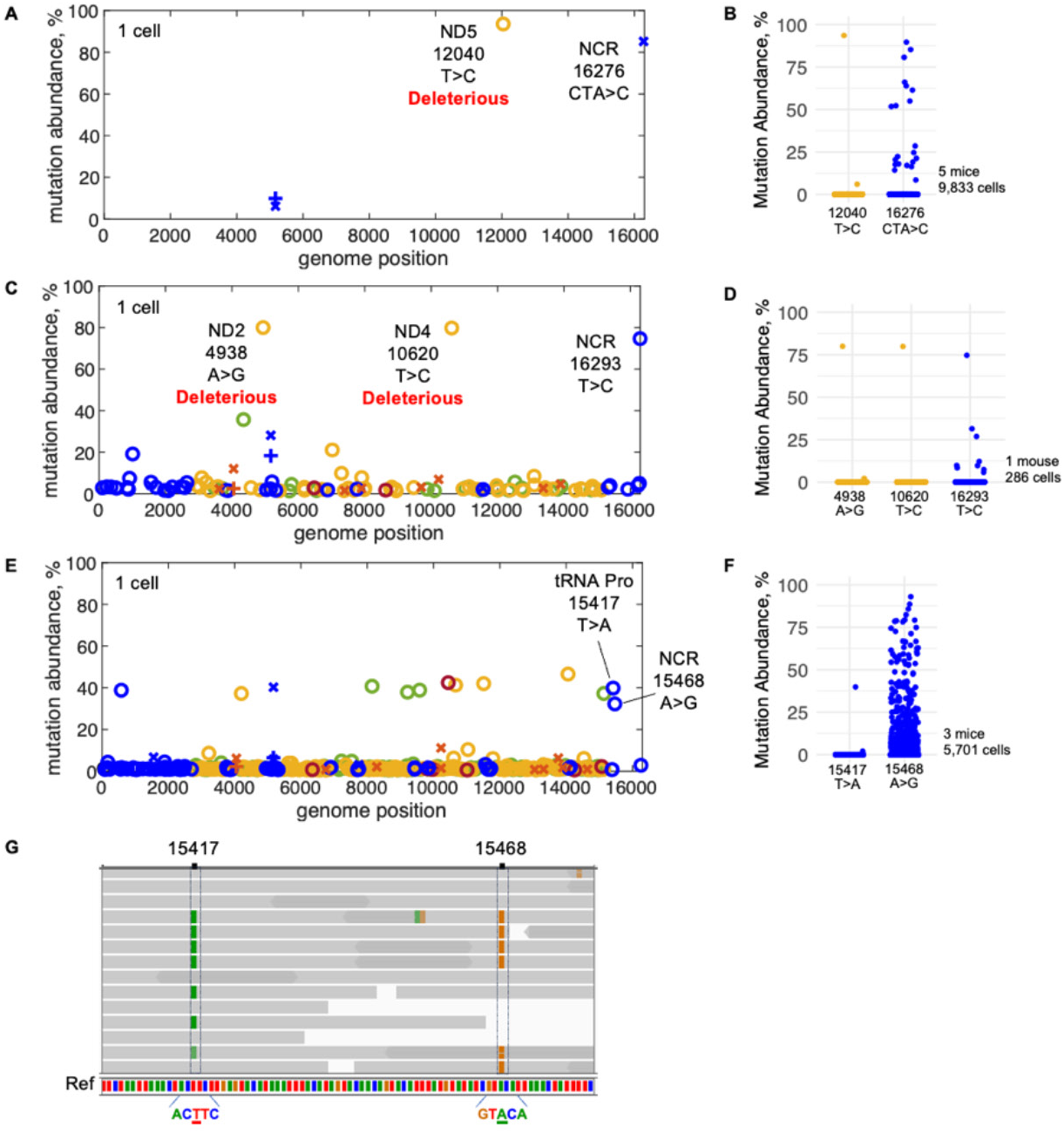
Passenger mutations piggyback on mitochondrial genomes with a competitive advantage. (A) A mtDNA mutation spectrum of a single liver cell from 24-month-old WT mouse showing two alleles at high abundance. (B) Abundance distribution of 12040T>C and 16276CTA>C mutations among all sequenced cells. (C) A mtDNA mutation spectrum of a single liver cell from 24-month-old heterozygous mutator mouse. (D) Abundance distribution of 4938A>G, 10620T>C and 16293T>C mutations among all sequenced cells of the sample. (E) A mtDNA mutation spectrum of a single liver cell from 24-month-old heterozygous mutator mouse. (F) Abundance distribution of 15417T>A and 15468A>G mutations among all sequenced cells. (G) Raw reads showing linkage of 15417T>A and 15468A>G mutations. Grey lines represent individual reads. Dark grey regions represent overlap of two opposing reads of the same DNA fragment. Green and orange bars mark mismatch between the read and reference sequences. See Figure 1C for the correspondence of symbols and allele type. Data in A, B, E, F and G were generated with 10X-based approach, data in C and D were generated with plate-based approach.

### Destructive selection

Since synonymous (S) mutations don’t interfere with protein function and nonsynonymous (NS) mutations can be disruptive, low ratios of accumulated NS to S mutations reflect elimination of NS alleles by purifying selection. Among closely related mammalian species, the mitochondrial coding sequences had an NS/S of 0.0588 indicating evolutionary conservation of protein function^44^. In contrast, mutant mtDNA alleles detected in aged mouse livers exhibited an average NS/S of 3.3, higher than expected even in the absence of purifying selection (Figure 6A). Similarly, the ratio of all detected nonsense (STOP) alleles to all detected S alleles was higher than expected (Figure 6B). These data suggest not only a lack of effective purifying selection as measured by the NS/S ratio of all detected alleles, but action of a selective force favoring NS mutations.

**Figure 6.**
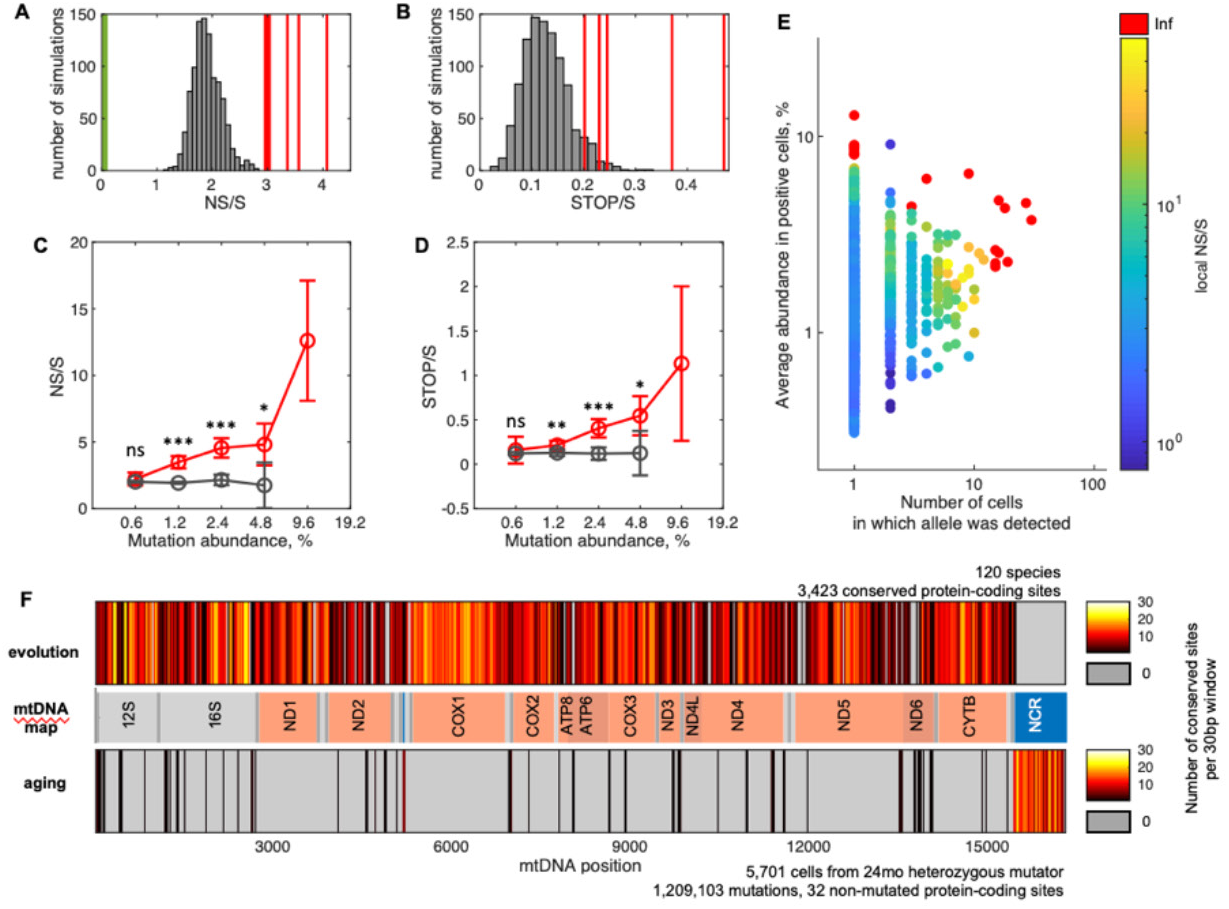
Excess of deleterious mtDNA mutations in aged mouse liver. (A, B) NS/S (A) and STOP/S (B) of all detected mutant alleles determined for six 24-month-old WT mouse livers (red lines) exceeds expected spread of NS/S and STOP/S (histograms) based on a 1,000 sets of simulations of mouse mtDNA random mutagenesis. The NS/S (0.0588) seen in evolution was taken from Pesole et al.^44^ (green line). Note that NS/S for three mice were very close and hence merged in a single thick line on the NS/S plot. Similarly, STOP/S ratios for two mice (0.2302 and 0.2299) are indistinguishable on the STOP/S plot. (C, D) NS mutations (C) and STOP mutations (D) selectively increase in abundance. Mean NS/S or STOP/S for mutations that fall in specified abundance intervals (log2 scale) from 6 mice (red line) or from 6 sets of 1,000 simulations of neutral behavior (grey line) with standard deviations. Model parameters: 10,000 genomes per cell, 3.16×10^-8^ mutation emergence rate, 80 generations. Mean values were computed when at least 3 out of 6 samples or simulations had finite NS/S ratios. Data within each abundance interval were tested against neutral model using two-sample t-test, ns – not significant, * - p<0.05, ** - p<0.01, *** - p<0.001. (E) Local NS/S for each NS allele on AAA vs C# plot shows clustering of NS alleles with similar ratios in 24-month-old WT mice. Data are the same as in Figure 3A, only NS alleles are plotted, N= 3 mice, 3,195 cells, 1,300 NS alleles. (F) Comparison of non-mutated (conserved) mtDNA sites in evolution (top bar) and in aging (bottom bar). Number of non-mutated sites within 30bp windows along mtDNA genome was plotted as a heatmap with yellow colors representing conserved regions and grey color marking windows in which all sites were changed in the data set. Middle bar represents mtDNA map. The list of species used for analysis of non-mutated sites in evolution is reported in Table S2 (n = 120). Aging data are from 24-month-old heterozygous mutator mice (n = 3 mice, 5,701 cells), which were also used in Figure 1E and S3. Data in this figure were generated with 10X-based approach.

When they first arise, mutations have not yet experienced selection, and their NS/S ought to approximate the expectation for random mutagenesis. NS mutations will directionally increase in abundance if they have a selective advantage. We divided our data into abundance categories and determined the NS/S ratio for the different groupings. For mutations present at low levels (0.3 – 0.6%), the NS/S ratio was close to random expectation (2.16 vs 1.9) but increased among mutations present at higher abundance. In contrast, simulated neutral mutations maintained a constant NS/S ratio across all abundance levels (Figure 6C). STOP/S also increased in the higher abundance bins (Figure 6D). This suggests that while mutations emerge randomly, NS and STOP mutations experience a selective advantage.

While clustering of positively selected NCR alleles at high abundance was obvious in AAA vs C# plots (Figure 3A), clustering of NS alleles was not. To better visualize the behavior of NS alleles on AAA vs C# plot we determined a local NS/S for every NS allele for three 24-month-old mice (see methods). As can be seen in Figure 6E, this approach showed clustering of high scoring NS alleles up and to the right of most NS alleles suggesting that select NS mutations are more strongly positively selected.

In contrast to our conclusion here, an increase in NS/S is often taken as an indication that a change in protein function has an advantage, often termed positive selection. Positive selection acts on rare alleles that improve fitness while mutant alleles reducing fitness will be subjected to negative/purifying selection. In contrast, if loss of function has an advantage, there should be no purifying selection for gene function. To obtain a genome wide view of the influence of function on selection, we compared the distribution of age-accumulated mutations across the genome to the distribution of changes occurring during evolution (Figure 6F). To illustrate a wide range in conservation, we divided the mouse mtDNA reference sequence into 30 base-pair windows and scored each window according to the number of base pairs that never change across a data set.

In a comparison of 120 distinct mammalian species, many windows have numerous conserved sites (white, yellow and orange-colored bands), and few windows (grey) in which all 30 base pairs change. In contrast, age-associated somatic mutations were widely distributed sparing few coding sites. The lack of evident conserved coding sequences in the aging data suggests a lack of selection for the function of coding sequences and is consistent with widespread destructive selection.

Notably, only the NCR region shows strong conservation during aging (Figure 6F). Because this region is involved in replication of the genome, we suggest that mutations at many sites in this region compromise replication and incur a competitive disadvantage while at a few NCR sites mutations are associated with the improved replicative competition of driver alleles.

### Selective forces in human liver

To test whether the same selective forces we observed in mice operate in human and how they play out on a longer time scale, we profiled mtDNA mutations in human hepatocytes from six de-identified individuals of known ages (Figures 7A and S6A). As expected for a long-lived organism (Figures 2C and 2D), many more mutations accumulated to higher levels in aged human samples than in mice (Figures 1C, 7A and S6A). This is especially true for the oldest (81-year-old) human sample (Figures S6A and S6B) in which many mutations have abundances near 100% (most of these are likely fixed and fall short of 100% abundance due to measurement inaccuracies; Figure S2 and methods). Two time-dependent mechanisms are expected to contribute to high abundance of alleles: random drift and positive selection (Figures S6C and S6D).

**Figure 7.**
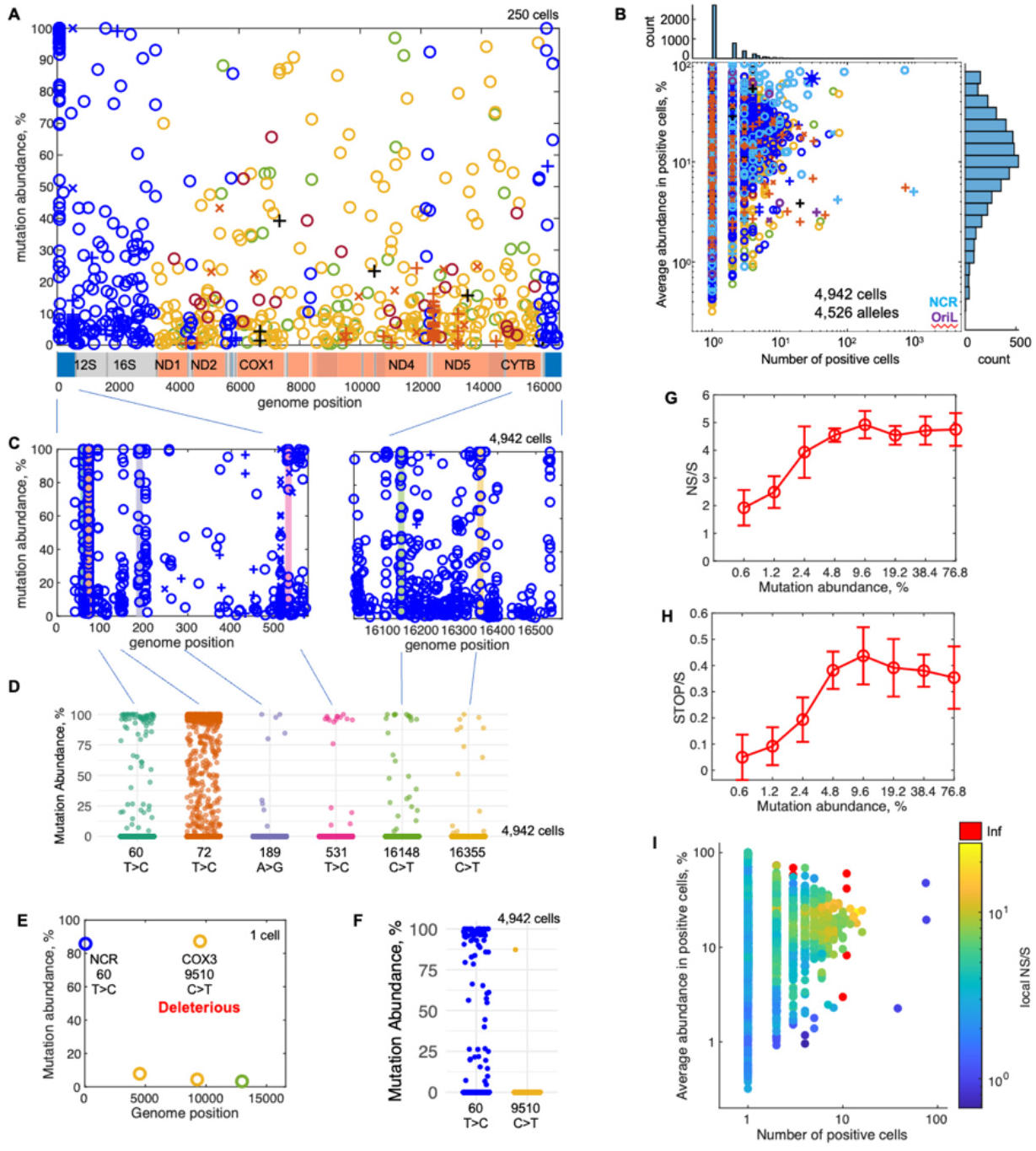
Selective forces impact competition among mtDNAs in human liver. (A) A spectrum of mtDNA mutations identified in 41-year-old human hepatocytes. Note that annotation of mouse and human mtDNAs differ with linearization of the human genome splitting the NCR in two. (B) AAA vs C# plot of 41-year-old human hepatocytes. Blue asterisk marks the 3243A>G allele. (C) A spectrum of mutations in the NCR of 4,942 hepatocytes from the 41-year-old human also shown in A. Colored bars indicate sites that meet our criteria for positively selected driver alleles. Mutations were classified as drivers if the allele was detected in at least 10 cells at levels above 50%, and there were more cells with >50% abundance than cells with <50% abundance. (D) Abundance-distribution of the driver mutations identified in NCR of 41-year-old human hepatocytes. (E) An example of a driver-passenger pair in a single liver cell from 41-year-old human. (F) Abundance distribution of the driver and the passenger alleles shown in (E) among the 4,942 sequenced cells of the sample. (G, H) The NS/S and STOP/S rises with increase in mutations abundance. (I) Local NS/S for each NS allele on AAA vs C# plot for 41-year-old human hepatocytes shown in (B). Data in this figure were generated with 10X-based approach.

As was the case with mouse data (Figure 3A), plotting the data in the AAA vs C# format shows enrichment of NCR alleles at high abundance (Figures 7B and S7), and the distribution of cellular abundance of these alleles in individual cells matches that expected for driver alleles (Figures 7A, 7C and 7D). Furthermore, examination of mutations in single cells shows evidence of driver/passenger linkage as we described in mouse (Figures 7E and 7F). We found variation in the drivers present in different individuals: in all we identified 18 driver alleles with high confidence among the six samples analyzed that lie within the NCR region. Several of these were previously identified as highly abundant alleles that occurred in a tissue-specific pattern in multiple individuals (Table S3)^19,30,45^. In accord with Samuels et al.^19^, we conclude that, a selfish/replicative advantage contributes to positive selection of NCR mutations in human liver as we saw in mouse. Furthermore, our findings agree with previous work in finding individual-to-individual variation in alleles exhibiting driver behavior (Table S3).

We next examined the data for signs of destructive selection. As in mice, the NS/S ratio increased among mutations present at higher abundance (Figure 7G). The STOP/S ratio behaved in a similar fashion (Figure 7H). We also examined the local NS/S ratios of all NS mutations alleles in AAA vs C# plots and again, as in mice, we found a cluster of NS alleles with an exceptionally high local NS/S ratio (Figure 7I). These data reveal that destructive selection operates in human liver in agreement with our data in mice and with a previous report^30^. Notably, over the lifetimes of the human samples, even a weak continuous positive selection drives alleles to fixation, yet the rise of the NS/S and STOP/S ratios reaches a plateau (Figures 7G and 7H) and NS alleles cluster at lower abundance levels than the driver alleles on AAA vs C# plots (Figures 7B and 7I). We suggest that the coefficient of selection imparted by destructive selection declines with increasing abundance, perhaps due to rising opposition from some form of purifying selection.

Mutations that enjoy a positive selection and yet are detrimental to function are expected to climb to unusual abundance in the human population and to make a dominant contribution to mitochondrial disease. Indeed, a well-known SNP (3243A>G) that is a common cause of mitochondrial disease satisfied our criteria for a driver allele in at least one individual (Figure 7B and Table S3) and showed a consistently high cellular abundance in our other human samples (Figure S7). Furthermore, the distributions of 3243A>G abundance in individual cells resemble those produced by inclusion of a positive selection coefficient (Figure S7E and S7F). The nature of this allele and its position on AAA vs C# plots (Figures S7A-S7D) could be consistent with either an especially strong destructive selection, and/or a replicative drive (see Discussion).

Together, these findings show that the same selective processes we identified in mice impact mtDNA in human hepatocytes and that time greatly magnifies the level of mutational accumulation.

## Discussion

Single cell sequencing of mtDNA from mammalian liver revealed how selection affects the accumulation of mitochondrial mutations with age. We expected that purifying selection would limit accumulation of mutations in two ways: mitochondrial quality control would target mitochondria carrying deleterious mutations for elimination, and death of metabolically compromised cells would eliminate cells with a heavy burden of mtDNA mutations. In contrast to a purifying action of selection, we found two pathways that accelerate the accumulation of mitochondrial mutations with age beyond that predicted for random mutation and neutral (non-selective) propagation, in mouse and in human hepatocytes. Here, we consider why these processes exist even though they appear to undermine fitness. We also discuss how they could impact age-associated accumulation of mtDNA mutations and influence the genetics of mitochondrial disease.

### Accumulation of mitochondrial mutations in the absence of selection is not so bad

Mutation-rate is a major determinant of how many mutant alleles emerge (Figure 1E), but it does not promote a rise in abundance of these alleles after they emerge. Even alleles with such high emergence rates that the mutation occurs multiple times in the same cell seldom rise to high cellular abundance (e.g., small indels in homopolymeric stretches; Figures 1C and 3A). Importantly, a cell can tolerate many mtDNA mutations at low abundance since other genomes can provide wild type function. Thus, mechanisms promoting a rise in the cellular abundance of mutations render these mutations impactful. Our single cell analysis of mtDNA mutations shows that only a few of the many possible alleles, those that benefit from positive selection, rise to high levels in hepatocytes. Consequently, we suggest that the types of selection we have characterized provides major routes to the emergence of cellular phenotypes, and hence will promote the progressive worsening of symptoms in patients with mitochondrial disease and enhance the deterioration of the mitochondrial genome with age.

While we conclude that neutral propagation of mtDNA alone cannot account for the observed age-dependent accumulation of mtDNA mutations in hepatocytes, we note that in other cell types, especially rapidly turning over cells with few copies of mtDNA, chance variation will be more impactful (Figure 2E). It will be important to understand the interaction of neutral variation with selective forces in various circumstances.

### A replicative advantage creates “selfish” selection

In mouse liver, we identified a prevalent class of mtDNA alleles that promote an increase in the relative abundance of affected genomes within the cells in which they are detected. Unlike other abundant mutations, such as maternally transmitted or early emerging mutations, these selfishly amplifying alleles appear in all mice in an inbred population, directionally increase in cellular abundance with age, and cluster in non-coding genomic sequences associated with replication. We refer to these alleles as driver mutations, because we find that they “drive” other linked mutations or “passengers” to high abundance.

We identified similar driver alleles in human hepatocytes. Since driver alleles achieve a high tissue abundance, it is not surprising that our list of human driver alleles overlaps with previously described mutations (Table S3). Notably, Samuels et al.^19^ described “recurrent, tissue-specific mutations” that occurred at high abundance in specific tissues of unrelated aged-individuals. Because these alleles were clustered in noncoding sequences adjacent to the origin of heavy strand replication and its regulatory sequences, it was argued that these mutations give genomes a replicative advantage in specific tissues. Our work further supports this and another past report^30^ in arguing for positive selection by a replicative advantage. This conclusion is in line with findings made in model systems from fungi to primates where mtDNAs have been found that preferentially replicate even when burdened with mutations deleterious to function^14–18^.

### Modulation of mitochondrial replicative drive by nuclear genes

Samuels et al.^19^ emphasized the tissue specificity of some members of his group of recurrent mutations. A similar tissue-specificity of competition between two mitochondrial genomes in a heteroplasmic mouse led Jenuth et al.^20^ to argue for “the existence of unknown, tissue-specific nuclear genes important in the interaction between the nuclear and mitochondrial genomes”. If differences in nuclear gene expression alters competition between mitochondrial genomes in different tissues, so might differences in the nuclear genome between individuals, a possibility supported by genetic demonstration of numerous nuclear modifiers of this competition in *Drosophila*^24^. Indeed, human drivers vary from individual to individual as expected for an influence of genetic background. For example, Samuels et al.^19^ found the 16093C>T allele at high levels in multiple tissues in only one of two individuals, and we saw this allele as a strong driver mutation in the hepatocytes of one individual but not in the hepatocytes of five others (Table S3). Similarly, other alleles show varied tissue distributions and sporadic variations from individual to individual^30,45^ (Table S3). Additionally, while we found the same driver alleles repeatedly in different individuals in an inbred strain of mice (C57BL6/J), among the sets of drivers we identified in two different mouse strains (C57BL6/J and mutator), only one driver was shared (Figure 4I).

Nuclear genes that promote replication of one mitochondrial genotype would disfavor other genotypes. Thus, a genome that is positively selected in one genetic background, can be negatively selected in another^24^. The extraordinary distinctions in the abundance of 16093C>T in different individuals is likely to include negative selection in individuals in which the allele was undetectable (Table S3). Finally, differences in nuclear gene expression associated with developmental stage, age, stress and diet are likely to alter the strength of selection for or against mtDNA loci sensitive to nuclear modification. Thus, while we have identified 18 driver alleles in human hepatocytes from six individuals (Table S3), a survey of additional tissues, individuals, and various life conditions is predicted to uncover many more.

### A nexus of genetic conflict

It is reasonable to assume that the NCR sequence controlling replication has been optimized over evolutionary time scales, in which case one would not expect to find driver alleles. However, evolution would select for the optimal replicator only in the germline, and somatic tissue-specific selection can “re-optimize” the NCR for replication in different somatic tissues. Since “re-optimization” of the NCR is the result of alterations of nuclear gene-action, change in the nuclear genome or changes in nuclear gene expression will give new driver alleles a selective advantage. Outbreeding, which changes the genetic background, will widely trigger selection to re-optimize mtDNA for replication, including in the germline. Driver alleles, while not detrimental in and of themselves, will amplify linked alleles, even if these passenger alleles are detrimental to the organism. Thus, by triggering selection for new drivers, outbreeding could promote germline trapping of passenger alleles, and such events could promote a heritable amplification of a disease allele. Given that such a process can be detrimental, we suggest that outbreeding-promoted selection for drivers might also be playing a role in the “hybrid breakdown” phenomenon which is suppressed by matching of maternal genotype and mtDNA^46,47^.

### Destructive selection

Previous studies have revealed an unexpected enrichment in NS alleles in mtDNA in the soma of *D. melanogaster*^33^, and humans^30^. Additionally, use of whole genome sequence data to identify population levels of mtDNA heteroplasmy also detected a selective rise in NS alleles in the elderly^48^. These studies suggest that alleles disrupting coding gene function have a selective advantage. We also see age-associated enrichment of NS alleles in mouse and human. The extent of the rise in abundance of different NS alleles varies, consistent with allele-specific impacts on gene function. But, given that such selection is ultimately destructive, why would such a process exist?

While there are several proposals for why defective mitochondrial genomes might have a selective advantage^33,49^, evidence from studies in *C. elegans* supports previous suggestions that use of feedback repression to control replication of mtDNA can give defective genomes an advantage^50–52^. According to these suggestions, mutations damaging function would allow the mutant genomes to avoid surveillance and the resulting feedback repression of their replication. Some strains of *C. elegans* harbor a large deletion mutant of mtDNA at high copy number^17^. Studies of genes impacting its copy number revealed the importance of ATFS-1. ATFS-1 is both a nuclear transcription factor regulating genes impacting the mitochondria and a mitochondrial protein where, among other things, it promotes mtDNA replication^53,54^. Functional mitochondrial genomes promote ATFS-1 degradation in the mitochondria, a negative-feedback on mtDNA replication that is not engaged by the deleted genome. It was found that mutational inactivation of this feedback reduced the copy number of the deleted genome relative to normal genomes, arguing that the defective genome obtained at least part of its advantage by evading this feedback loop. Based on these findings, we suggest that evolutionary fitness benefits of feedback repression to control mtDNA copy number in the soma have outweighed the costs incurred by promoting accumulation of defective mitochondrial genomes.

Whatever its mechanism, destructive selection acts widely to increase the overall NS/S ratio of somatic mtDNA mutations. Its actions, if unopposed by other selective actions, would promote a relentless rise in mutations that are deleterious to function within individual cells in which such mutations occur.

### Is there a covert version of purifying selection?

Purifying selection is a form of negative selection that eliminates mutations compromising fitness of the host organism. Its action during evolution results in conservation of important coding sequences. Our comparison of sequences conserved in evolution to sequences conserved in hepatocytes during mouse aging detected no obvious signature of purifying selection acting on coding sequences during aging (Figure 6F). Furthermore, we found that in mice the NS/S ratio increased progressively with the abundance of the surveyed alleles, arguing for continuous destructive selection at least in the early stages of accumulation. Despite these findings, it remains possible that purifying selection also occurs and is beneficial. Destructive selection acts in opposition to purifying selection. Consequently, if both operate, the dominant one will prevail, thus masking the other in our analysis of the net outcome.

*C. elegans* offers a view of the interplay of destructive selection and purifying selection and how this genetic conflict plays out at the whole animal level^55^. When propagated as small populations, *C. elegans* lineages accumulate high levels of deleted mtDNAs despite costs to fitness. These lineages persist with a relatively stable mix of intact and deleted genomes, apparently by balancing a destructive selection promoting amplification of the deleted genome by a fitness based purifying selection that continuously culls individuals with an especially high burden of the defective genome. We suggest that a process parallel to this occurs in the soma in mammals. We propose that, as detrimental alleles increase in cellular abundance, they compromise cell fitness leading to growth arrest or cell elimination, resulting in a counteracting purifying selection that stalls the climb in abundance of deleterious mutations. Our data suggest that this “flipping point” occurs at a relatively high abundance of the mutant alleles when they compromise cell function. The balance of destructive selection and purifying selection, whether in the case of *C. elegans* deletions or the proposed balance in somatic tissues, comes at a fitness cost — either organisms or cells are eliminated. Accordingly, destructive selection is, at its root, a detrimental influence.

The conservation of sequence in the NCR region seen during aging is likely due to a different kind of negative selection. Mutations in this region are likely to compromise replication and suffer a strong competitive disadvantage. Only rare alleles will improve replication to generate a “driver” with a selective advantage. Accordingly, sequences in this region are likely to be under especially strong selective pressures, whether negative or positive.

### Selection could promote inheritance and accumulation of mtDNA disease alleles

The prevailing class of inherited mtDNA disease alleles are transmitted in combination with wild-type genomes. Such heteroplasmic alleles can circulate in the population without clinically identified symptoms and can sporadically rise in abundance to levels that produce disease. Even when the disease allele is detected, unpredictable shifts in abundance and varied tissue distributions bedevil accurate prognoses of transmission and of disease severity. A better understanding of the contributions of selection to disease allele propagation is likely to improve clinical management and perhaps lead to the discovery of approaches to limit disease allele levels in the population and disease severity in affected individuals.

There are many mtDNA-associated mitochondrial disease alleles, but one, 3243A>G, is responsible for more cases of mitochondrial disease than any other identified mtDNA single nucleotide change^56^. The 3243A>G mutation disrupts the gene TM-TL1 which encodes tRNA-leu(UUR)^57^, as well as altering a sequence required for binding of a transcriptional terminator^58^. The mutation reduces mitochondrial translation^59^, which is thought to be responsible for the deleterious consequences of the allele. However, other mutant alleles share these molecular defects yet lack the high incidence of 3243A>G mutation, leaving us without an explanation for its disproportionate prevalence.

There have been many investigations of 3243A>G abundance in patients and their relatives. In these cases, affected individuals inherited multiple copies of the mutant genome which were present in a large fraction of their cells. Our study is unusual in examining behavior of this mutation as it emerges in somatic tissues *de novo*. In total, we analyzed 25,997 hepatocytes from 6 individuals and detected the 3243A>G allele in 72 cells (∼0.3%). All samples included cells with an unusually high accumulation indicating positive selection of the 3243A>G allele (Figure S7), which is consistent with a previous observation^51^. Despite an origin from a somatic mutation, 3243A>G on average reached 50% abundance in these rare positive cells which is close to the upper range reported for patients with severe mitochondrial disease and translates into a staggering ∼2,500-fold increase in abundance from the initiating event. We propose that the positive selection that we see following somatic emergence of this mutation contributes to both the population prevalence of 3243A>G allele as well as to disease progression in affected individuals.

Studies of 3243A>G in patients show particularly high levels of the allele in muscle (e.g 77%^28^), and especially low levels in peripheral blood mononuclear cells. Extensive analyses of 3243A>G in patient peripheral blood have shown an age dependent decline suggesting purifying/negative selection particularly in the lymphoid lineage^26–28^. A detailed recent study using single cell analyses suggests that purifying selection in the lymphoid lineage is not limited to 3243A>G allele as the levels of mtDNA carrying large deletions is also reduced with patient’s age in these cells^60^. While these studies reveal complexities in the selective events influencing mitochondrial mutations in different cell types, we suggest that, in at least some cell types, newly emerged 3243G>A mutations are carried to high cellular abundance by positive selection as we have seen in hepatocytes. Importantly, even if it were only to occur in some genetic backgrounds, positive selection of 3243A>G in the germline^61^ could account for its prevalence and differential selection in different tissues could account for the diversity of disease presentations.

### Limitations of this study

While our study argues strongly that the net outcome of the actions of selection in the livers of mice and humans promotes the accumulation of mutant mtDNAs beyond what is expected from neutral models, there have been several reports of the potential for purifying selection to do the opposite^62^. There is evidence that cell death and/or reduced proliferation due to compromised mitochondrial function can select against cells with a high burden of dysfunctional mitochondrial genomes^29,60^, and that quality control eliminates defective mitochondria either by autophagy^25,63^ or mitocytosis^64^, or selectively promotes biogenesis of functional mitochondria^3,5^. Importantly, the net outcome we report reveals the dominant selection, leaving open the possibility that purifying selection has a modulating action in liver, and perhaps a dominating influence in other tissues or developmental stages. Both possibilities could be explored by assessing whether mutations compromising quality control mechanisms impact mtDNA integrity in liver and other tissues during aging.

Unfortunately, detailing the accumulation of mutations with age highlights some interesting “why” questions without answering them. Is the extraordinary germline conservation of mtDNA entirely due to selection for fitness, or do quality control filters influence mammalian transmission of mtDNA mutations? And given the existence of quality control, why doesn’t it effectively safeguard mtDNA during aging? Might it be that evolutionary pressure for elite performance in the adult is not compatible with conditions needed for quality control of mtDNA genes?

## Acknowledgments

We are grateful to Saul Villeda for sharing aged mice and support with establishment of mutator mouse colony. We thank UCSF LARC for help with mouse husbandry. We thank Spyros Darmanis (CZ Biohub) for sharing index primer sequences for mtATAC library preparation. We thank Eric Chow and UCSF CAT for providing access to basic and cutting-edge equipment, support and advice, and Steven Deluca for the suggestion to use ATAC-seq for mtDNA profiling. Sequencing was performed at the UCSF CAT, supported by UCSF PBBR, RRP IMIA, and NIH 1S10OD028511-01 grants. This study was supported in part by the Liver Cell Isolation, Analysis & Immunology Core of the UCSF Liver Center (P30DK026743) and HDFCCC Laboratory for Cell Analysis Shared Resource Facility through a grant from NIH (P30CA082103). Portions of this work were performed on the Wynton HPC Co-Op cluster which is supported by UCSF research faculty and UCSF institutional funds. We thank the UCSF Wynton team for their ongoing technical support of the Wynton environment. This work was funded by Larry L. Hillblom Foundation (2018-A-028-FEL to E.K., 2019-A-011-NET to P.OF. and 2019 John S. Spice award in Aging to E.K.), UCSF Program for Breakthrough Biomedical Research (2019-2020 New Frontier Research Award to P.OF and Saul Villeda and 2021-2022 Postdoc Independent Research Grant to E.K.), CNV Stiftung to E.K., NIH R35GM136324 to P.OF. and NIH R33CA247744 to Z.J.G. We thank Sandy Johnson for critical reading of the manuscript.

## Author contributions

E.K. and P.H.O’F. conceived the project, designed experiments, interpreted the results, and secured funding. E.K. and D.N.C. adapted MULTI-ATAC and 10X scATAC for profiling mtDNA sequences and performed the initial set of experiments employing the 10X-based approach. E.K. performed all other experiments and data analysis. Z.J.G. provided expertise for MULTI-ATAC and 10X scATAC. E.K and P.H.O’F. wrote the manuscript with input from all authors.

## Declaration of interests

Z.J.G. is an author on a patent on MULTI-seq technology, and it has been licensed to Millipore.

## Supplemental information

Document S1. Figures S1–S7 and legends.

Tables S1-S3. Excel files containing additional data too large to fit in a PDF.

## Material and Methods

### Animals

C57BL6/J, mtPWD (C57BL/6J-mt^PWD/Ph^/ForeJ) and mutator (B6.129S7(Cg)-Polg^tm1Prol^/J) mice were obtained from The Jackson Laboratory. Heterozygous mutator males were bred with C57BL6/J females to produce heterozygous progeny for experiments. Mice were housed in a specific pathogen-free facility with a standard 12-h light/dark cycle at the University of California, San Francisco, and given food and water *ad libitum*. Experiments were conducted in accordance with institutional guidelines approved by the University of California, San Francisco Institutional Animal Care and Use Committee.

### ddPCR

Genomic DNA was isolated from 25mg of liver tissue using DNeasy Blood and Tissue kit (Qiagen, 69506) according to manufacturer’s guidelines. Primers and probes were synthesized by Integrated DNA Technologies (IDT) and their sequences are provided in Table S1. WT C57BL6/J mtDNA sequence (p15196 – p136) was cloned into pGEM-T (Promega, A1360) vector and used as pure WT control. A 500bp fragment of mouse mtDNA containing 15468A>G or 16012G>A mutations was synthesized by IDT and cloned in pUCIDT-AMP vector to use as positive controls. The ddPCR reaction mixture contained ddPCR Super Mix for Probes (Bio-Rad, 1863024), 900 nM of forward primer, 900 nM of reverse primer, 250 nM of WT probe, 250 nM of mutant probe, 0.5 μL of restriction enzyme (HaeIII; NEB, R0108L) and template DNA. Template DNA concentration was adjusted to be below 3,500 mtDNA copies per microliter of ddPCR reaction mixture. 20 μL of the reaction mixture and 70 μL of oil (Bio-Rad, 1863005) were loaded on a DG8 cartridge (Bio-Rad, 1864007) for droplet generation on QX100 Droplet Generator (Bio-Rad). 40 μL of droplet emulsion were transferred to 96-well plate (Bio-Rad, 12001925) and sealed with a pierceable foil (Bio-Rad, 1814040) using PX1 PCR plate sealer (Bio-Rad). The optimized PCR thermal cycling was conducted on a conventional PCR machine (Bio-Rad, C1000 Touch). Thermocycling conditions for the 15468A>G assay: 10 min polymerase activation at 95°C; 40 cycles of denaturation at 94°C for 30 s, ramp rate 1°C/s, and combined annealing-extension at 54°C for 2 min, ramp rate 1°C/s; incubation at 98°C for 10 min. Thermocycling condition for the 16012G>A assay: 10 min polymerase activation at 95°C; 45 cycles of denaturation at 94 °C for 30 s, ramp rate 1°C/s, and combined annealing-extension at 52°C for 2 min, ramp rate 1°C/s; incubation at 98°C for 10 min. After thermocycling, samples were cooled to room temperature and analyzed on the QX100/200 Droplet Reader (Bio-Rad).

Results were analyzed with QuantaSoft Analysis Pro v.1.0.596 software (Bio-Rad).

### Hepatocytes isolation

Mouse hepatocytes were isolated by a two-step perfusion technique. Briefly, mouse was anesthetized by isoflurane (Piramal Critical Care). Mouse liver and heart were exposed by opening the abdomen and cutting the diaphragm away. The portal vein was cut and immediately the *inferior vena cava* was cannulated via the right atrium with a 22-gauge catheter (Exel International, 26746). Liver was perfused with liver perfusion medium (Gibco, 17701038) for 3 minutes and then with liver digest medium (Gibco, 17703034) for 7 minutes using a peristaltic pump (Gilson, Minipuls 3). Pump was set to 4.4 ml per minute and solutions were kept at 37°C. After perfusion the liver was dissected out, placed in a petri dish with hepatocyte plating medium (DME H21 [high glucose, UCSF Cell Culture Facility # CCFAA005-066R02] supplemented with 1x PenStrep solution [UCSF Cell Culture Facility # CCFGK004-066M02], 1x Insulin-Transferrin-Selenium solution [GIBCO #41400-045] and 5% Fetal Bovine Serum [UCSF Cell Culture Facility # CCFAP002-061J02]) and cut into small pieces. Liver fragments were passed through a sterile piece of gauze. Hepatocytes were separated from non-parenchymal cells by centrifugation through 50% isotonic Percoll (Fisher# NC9256155) solution in HAMS/DMEM (1 packet Hams F12 [GIBCO # 21700-075], 1 packet DMEM [GIBCO # 12800-017], 4.875g sodium bicarbonate, 20mL of a 1M HEPES pH 7.4, 20mL of a 100X Pen/Strep solution, 2L H2O) at 169g for 15 min. Isolated hepatocytes were used immediately for FACS or frozen in BAMBANKER (GC LYMPHOTEC, CS-02-001) and stored at −80°C for future experiments.

Cryopreserved deidentified human hepatocytes were purchased from Xenotech, Lonza or UCSF Liver Center.

### Cell sorting

Isolated hepatocytes were resuspended in PBS, stained with 5 μg/mL propidium iodide (Invitrogen, P1304MP) to mark dead cells, and kept on ice until FACS. Right before sorting hepatocytes were strained through 35-40 um cell strainer. Sorting was performed on FACSAriaII (Becton Dickinson) using 100 um nozzle. Instrument was calibrated using 23.9 um beads (Spherotech, ACURFP2.5-250-5). Single hepatocytes were sorted into 384-well plates (Bio-Rad, HSP3801 or 4titude, 4ti-0384) containing 0.45 ul of TD buffer (10 mM TrisHCl pH 8.0, 5 mM MgCl_2_, 10% dimethylformamide). Due to their large size and extreme size variability, sorting of mouse hepatocytes was inefficient and only 40-60% of wells contained cells while the rest of the wells were empty. Human hepatocytes sorting efficiency was 90%. One column of a plate was left empty to serve as a negative control. Immediately after sorting plates with hepatocytes were sealed with foil (Bio-Rad, MSF1001 or 4titude, 4ti-0500FL), briefly centrifuged, frozen on dry ice and stored at −80°C.

### Single cell ddPCR

mtDNA copy number was quantified using single cell ddPCR. Frozen 384-well plate with single hepatocytes in 0.45 ul of TD buffer was thawed on ice. To lyse hepatocytes 0.45 ul of solution containing 10mM Tris-HCl pH 8.0, 50mM NaCl, 40 ng/uL MS2 RNA, 0.4% SDS and proteinase K (8U/ml; NEB, P8107S) was added to 96 wells of the plate with help of acoustic liquid handler Echo 525 (Beckman Coulter). Wells without cells were used to prepare positive and negative controls. gBlock encompassing amplified sequence was synthesized by IDT and used as a positive control. After lysis and control solutions were added, the plate was sealed, briefly centrifuged, and incubated at 50°C for 15 min and then at 95°C for 10 min. After lysis, ddPCR master mix (ddPCR Super Mix for Probes (Bio-Rad, 1863024), 250 nM of forward primer, 250 nM of reverse primer, 250 nM of probe, 0.5 μL of restriction enzyme [AluI (NEB, R0137L) for mouse assay and HaeIII (NEB, R0108L) for human assay]) was added to each of 96 wells, plate was sealed and vigorously vortexed, briefly centrifuged and incubated at 37°C for 15 min to digest DNA. After restriction enzyme digestion, the plate was vigorously vortexed, briefly centrifuged and 20 ul of the reaction mixture was used for ddPCR as described above. Primers and probes sequences are provided in Table S1. Thermocycling conditions for mouse mtDNA copy number assay: 10 min polymerase activation at 95°C; 40 cycles of denaturation at 94°C for 30 sec, ramp rate 2°C/s, and combined annealing-extension at 52°C for 1 min, ramp rate 2°C/s; incubation at 98°C for 10 min. Thermocycling conditions for human mtDNA copy number assay: 10 min polymerase activation at 95°C; 40 cycles of denaturation at 94°C for 30 sec, ramp rate 2°C/s, and combined annealing-extension at 56°C for 1 min, ramp rate 2°C/s; incubation at 98°C for 10 min.

### Plate-based single cell mtATAC

Frozen 384-well plates with single hepatocytes were thawed on ice. Hepatocytes were lysed and DNA was tagmented in a single step. To this end, 0.45 ul of lysis solution (1% n-Dodecyl β-D-maltoside [final concentration 0.5%], 90 mM NaCl [final concentration 45 mM], 10mM TrisHCl pH 8.0, 5 mM MgCl_2_, 10% dimethylformamide) supplemented with Tn5 (Illumina, 20034197; 1.5 ul of enzyme for 150 ul of lysis solution) was added to each well of a plate with help of Echo 525. Then, plates were sealed with foil (Bio-Rad, MSB1001), briefly centrifuged, and incubated at 37°C for 30 min. After lysis and tagmentation, Tn5 was stripped off DNA. To this end, 0.1 ul of 2% SDS was added to each well of a plate (final concentration 0.2%) using Echo 525, plates were sealed with a foil, briefly centrifuged, and incubated at 65°C for 15 min. Next, mtATAC libraries were constructed by PCR amplification of DNA fragments created by Tn5 with unique dual index primers for each well of a plate. To this end, PCR master mix (NEB, M0544S), tween-20 (final concentration 0.34%) and unique dual index primers (final concentration 500 nM; sequences are provided in Table S1; IDT) were added to each well of the plate using Echo 525 (final volume 3 ul), plate was sealed with a foil, briefly centrifuged and thermocycled as follows: incubation at 72°C for 5 min to fill the gaps; initial denaturation at 98°C for 30 s; 16 cycles of denaturation at 98°C 10 sec and combined annealing-extension at 65°C 75 sec; final extension at 65°C 5 min. Incubation and PCR were performed in a standard thermocycler (Bio-Rad, C1000 Touch or S1000). Uniquely labeled libraries from one or several plates were pooled together at equal volumes and cleaned up using home-made SPRI beads twice^1^. The first cleanup was one-sided with 1.2 beads to library volume ratio. The second cleanup was two-sided with 0.5 ratio followed by 1.2 ratio. Cleaned up libraries were eluted in 20 ul of TE buffer. To quality control and quantify libraries 1 ul of cleaned mtATAC library was run on Bioanalyzer (Agilent). mtATAC libraries were sequenced on MiSeq (Illumina) using MiSeq Reagent Kit v2, 300-cycles (Illumina, MS-102-2002) as 151×12×12×151.

### 10X-based single cell mtATAC

Frozen hepatocytes were thawed, washed with PBS (Gibco, 10010-023) and fixed in 1% PFA for 10 min at RT. After fixation PFA was quenched with glycine (125 mM final concentration) and washed with cold PBS supplemented with 1% BSA (Sigma, A1953). Next, hepatocytes were permeabilized. To this end, 1 million fixed cells were resuspended in 200 ul of lysis solution (0.5% n-Dodecyl β-D-maltoside, 45 mM NaCl, 10 mM Tris-HCl pH 8.0, 5 mM MgCl_2_, 10% dimethylformamide) and incubated on ice for 5 min. For human hepatocytes n-Dodecyl β-D-maltoside concentration was reduced to 0.1%. Permeabilization was stopped by adding 1.8 ml of wash buffer (45 mM NaCl, 10 mM Tris-HCl pH 8.0, 5 mM MgCl_2,_ 1% BSA). To enable pooling of distinct samples in a single 10X experiment, permeabilized cells from different mice were labeled with unique DNA barcode complexes (MULTI-ATAC; Conrad et al., in preparation).

MULTI-ATAC barcoding was also performed when cells from a single individual were analyzed to improve identification of multiplets. In this case, an individual sample was divided into 3 to 7 fractions and each fraction was labeled with a unique MULTI-ATAC barcode. To this end, Lignoceric Anchor oligo (2 uM; Sigma, LMO001A) was mixed with a unique barcode oligo (1 uM; BC) and reverse primer (1 uM; BE) at 1:1:1 molar ratio to form Anchor-BC-BE Complex (20x, 1 uM). Note that BC contains an 8-nucleotide-long stretch of random nucleotides to serve as a unique molecular identifier (UMI) to enable barcode counting. Permeabilized hepatocytes were resuspended at 10^6^ cells/mL in cold PBS and Anchor-BC-BE Complex was added to cell suspension (final 1x, 50 nM) followed by incubation on ice for 5 min. To stabilize labeling, Palmitic Co-anchor oligo (2uM, 20x; Sigma, LMO001B) was added to the cell suspension (final 1x, 50 nM) followed by additional incubation on ice for 5 min. After MULTI-ATAC barcoding, unbound complexes were washed away with PBS supplemented with 2% BSA, hepatocytes isolated from different individuals were pooled together, resuspended in diluted nuclei buffer, passed through 35-40um cell strainer, and used to prepare 10X-mtATAC libraries using Chromium Next GEM Single Cell ATAC Reagent kit (10X Genomics, PN-1000176 and PN-1000406) according to the manufactures protocol (CG000209 Rev F and CG000496 Rev B) with 2 minor modifications. First, after step 3.2o 1 ul of the sample was used to prepare the MULTI-ATAC barcode library (described below). Second, the remaining 39 ul were used in step 4.1 where SI-PCR Primer B concentration was increased to 100 uM. Before permeabilization, after permeabilization and after MULTI-ATAC barcoding cells were pelleted by centrifugation at 100g for 3 min, 300g for 3 min and 500g for 5 min, respectively.

To prepare MULTI-ATAC barcode libraries, 1ul of sample from 3.2o step was amplified in a PCR reaction: 1 ul of sample, 500nM SI-PCR-B primer, 500nM TruSeq primer, 1x Kapa HiFi HotStart ReadyMix (Roche, KK2601). The reaction mixture was thermocycled using the following conditions: 5 min polymerase activation at 95°C; 14 cycles of denaturation at 98°C for 20 sec, annealing at 67°C for 30 sec and extension at 72°C for 20 sec; incubation at 72°C for 1 min.

To quality control and quantify libraries, 1 ul of 1:5 diluted 10X-mtATAC and MULTI-ATAC barcode libraries were run on Bioanalyzer. 10X-mtATAC and MULTI-ATAC barcode libraries were pooled together and sequenced on NovaSeq6000, S1 200 as 101×12×24×101 or NovaSeq X, 10B as 51×12×24×51 or 151×12×24×151. For optimal demultiplexing we aimed to obtain 5,000 MULTI-ATAC barcode reads per cell.

This method is prone to low level leakage of mutation signal between cells (Figure S2). Since inbred mice have identical mitochondrial genomes and mutations are very rare this leakage becomes noticeable only if clonal mutations are present. Unlike inbred mice, humans have multiple haplotype- and individual-specific mtDNA variants. Consequently, leakage is noticeable at multiple sites if different human samples are mixed in a single experiment.

Therefore, to simplify downstream analysis all human samples were processed individually. Importantly, leakage also affects our readings of fixed mutations: as predominant WT signal leaks into cells with fixed mutations, often times we detect these mutations at levels just below 100% instead of 100%.

### Sequencing data analysis

#### Reads mapping, coverage and variant analysis

The nucleus contains multiple segments derived from mtDNA sequence, so called NUMTs. When sequencing reads from ATAC experiments are aligned to the whole genome a lot of truly mitochondrial reads are erroneously mapped to the NUMTs. To avoid incorrect mapping of mitochondrial reads to the nuclear genome sequencing reads from the plate-based approach were aligned directly to the mitochondrial genome. Specifically, reads were aligned to the mouse mitochondrial genome (NC_005089) with *bwa*^2^ (v0.7.17) *using* the *BWA*-*MEM* algorithm. Samples with less than 10,000 reads mapping to mtDNA (chrM) were excluded from further analysis. Duplicate reads were marked with *Picard tool*^3^ (v2.27.4). Mapped reads were filtered with *bamtools*^4^ (v2.5.2; filter-mapQuality “>=20”-isPaired “true”-isProperPair “true”). Coverage was determined with *samtools depth*^5^ (v1.16.1). SNPs and small indels were called using *Freebayes*^6^ (v1.3.6;-C 5-F 0.003-p 1--pooled-discrete--pooled-continuous -m 30 -q 30 - -min-coverage 10). Multiallelic sites were split into multiple rows using *bcftools*^5^ (v1.16; norm - Ov m-both). Variants were filtered using *vcffilter*^7^ (vcflib v1.0.3;-f “SAF >1”-f “SAR >1”). Complex alleles were reduced to primitive alleles using *vcfallelicprimitives* and sorted with *vcfstreamsort*^7^ (vcflib v1.0.3). This process occasionally created duplicate variants where mutation counts were split between the records which led to incorrect mutation frequency calculation. This issue was fixed by merging duplicated records in a single entry with alternative allele counts summed together. This was done after vfc files from individual cells were merged using *bcftools* (merge-m none) and the resulting vcf file was converted to tab delimited file using *vcf2tsv*^7^ (vcflib v1.0.3). The variants were spot checked in IGV^8,9^ (v 2.4.16). Variants annotation (synonymous, non-synonymous, stop-gain and etc.) was done using *SnpEff*^10^ (v5.0). When the same variant had multiple annotation (e.g., due to overlap of protein coding sequences) the most severe annotation was used. *Pindel*^11^ (0.2.5b9) was used to detect large-scale deletions (minimum deletion size 10bp, at least 10 supports). *SIFT*^12^ was used to predict whether mutation affects protein function.

Sequencing data from 10X-based experiments were first processed with *Cell Ranger ATAC* (10X, v 2.1.0) using blacklisted reference genomes^13^. Blacklisting was necessary to prevent erroneous mapping of mtDNA fragments to the nuclear DNA. Because our samples are non-standard and predominantly contain mtDNA reads, *Cell Ranger ATAC* does not discriminate well between empty droplets and droplets containing cells. To classify droplets into those containing cells and those that are empty as well as to assign cells to samples in multiplexed experiments and identify droplets with multiple cells, we relied on MULTI-ATAC barcode UMI counts. Barcode UMI counts have a bimodal distribution where positive (high-count) and negative (low-count) droplets for a specific barcode are clearly separated. To this end, a list of droplets that contained at least 1,000 or more reads that passed filters (metrics provided by *Cell Ranger ATAC*; the cutoff was set to include 1.5 to 2 times more droplets than expected recovery) along with MULTI-ATAC barcode library FASTQ files were supplied to *deMULTIplex2*^14^. For proper performance, *deMULTIplex2* requires removal of most empty droplets before demultiplexing. Hence, the barcodes count matrix created by deMULTIplex2 was filtered based on total number of MULTI-ATAC barcode UMIs. The cutoff was determined by plotting a histogram of barcodes counts and finding the middle between the two peaks representing positive and negative droplets. Whenever possible faithfulness of demultiplexing was controlled by analysis of distribution of sample-specific SNPs among multiplexed samples. Reads from droplets that were classified as carrying a single cell were subset from *possorted_bam.bam* file generated by *Cell Ranger ATAC* into separate bam files using *samtools*^5^ (v1.16.1). Reads deduplication, reads filtering, variant calling, variant filtering and variant annotation were the same as for plate-based approach. Cells with average mtDNA (chrM) coverage less than 50 were excluded from the analysis.

In addition to standard variant filtering the following calls were excluded from the final datasets. Large-scale deletions in minor and major arcs of mouse mtDNA cause misalignment at the imperfect repeat regions creating false 4920C>T and 4925C>G, and 8686T>C, 14251T>A and 14260T>G mutations, respectively, that were excluded from the final dataset. Due to high number of mismatches between C57BL6/J mtDNA (NC_005089) and PWD mtDNA (DQ874614) sequences in NCR (p15400 - p15600), reads from mtPWD samples do not map to C57BL6/J reference in this region. Therefore, PWD-specific SNPs in this region were excluded from the dataset when analyzing C57BL6/J and mtPWD mixing results. Human mtDNA reference sequence (NC_012920) contains N at position 3107 which denotes deletion. This N is misinterpreted by the aligner as any nucleotide which leads to 3107N>C, 3107N>T and 3109T>C false mutations. These calls were excluded from the final dataset. The region between p300 and p320 of human mtDNA had low coverage and multiple sequencing errors making it difficult to distinguish true and false mutations. Therefore, variants in this region were removed from the dataset. Finally, variants with mean abundance above 90% in a human sample were considered haplotype or individual-specific polymorphisms and were excluded from the list of mutations.

Despite our best effort to remove false calls from the dataset, there are some remaining artifacts present, in particular sequencing or alignment errors at the ends of the reads. Those usually are present at very low levels and are unlikely to have any impact on our conclusions. All the specific mutations (such as driver and passenger alleles) that we rely on to draw conclusions were hand checked in IGV.

#### Percent of reads mapping to mtDNA

Total number of reads and number of reads mapping to mtDNA (chrM) were calculated using *samtools view*. Number of reads mapped to mtDNA was divided by total number of reads and the values were converted into %.

#### Number of mutant alleles

To calculate number of mutant alleles (Figure S3E) we used data produced with 10X-based approach. The number of unique mutant alleles strongly depends on coverage and number of analyzed cells. To mitigate biases due to coverage differences between samples we subsampled deduplicated and filtered bam files for individual cells to 100,000 mtDNA (chrM) mapped reads. Cells that had fewer reads were excluded from the analysis. Subsampled files were used for variant calling as described above. Finally, we normalized samples by analyzing equal number of cells from each sample.

#### NS/S and STOP/S

NS/S and STOP/S were calculated using three approaches. In the first approach, ratios were calculated for all detected alleles in a sample, so that each unique allele was counted only once (Figures 6A and 6B).

In the second approach, ratios were calculated for all SNP mutations within abundance intervals (specified in the figure; Figures 6C, 6D, 7G and 7H), i.e. each allele was scored according to the number of cells in which it was detected. In this case clonal mutations were excluded from the analysis as these mutations have a major impact on the ratio but unlikely to reflect impact of selective forces. Clonal mutations are expected to be present in unusually high number of cells in only one or few of the examined individuals. To identify such alleles, we applied Kruskal-Wallis test to the mutation cellular abundance data from multiple individuals. Bonferroni adjustment of the p-values was performed to account for the large number of tested alleles. Adjusted p-value of 0.05 was used as a cutoff for decision between clonal and non-clonal alleles. This method is sensitive to difference in coverage between analyzed samples, therefore it was applied to mouse data from different experiments separately as those tend to have different coverage. In case of human data, where each sample was prepared individually and hence coverage varies among all the samples, we tried two approaches. In the first approach, we subsampled human data to 100,000 mtDNA mapped reads per cell and then applied Kruskal-Wallis test. The obtained list of clonal alleles was then used to remove clones from the original dataset. This approach might miss clonal mutations with low cellular abundance. In the second approach we applied the test directly to the dataset without adjustment for coverage. In this case along with true clonal mutations and number of low abundance alleles were removed. While both approaches identified different number of clonal alleles, the final results (shape of the curve in Figures 7G and 7H) were similar. Data presented in Figures 7G and 7H were generated with the first approach.

In the third approach, we calculated local NS/S ratio on an AAA vs C# plot (Figures 6E and 7I). For each NS allele on log10 transformed AAA vs C# plot we counted number of NS and S alleles within a circle centered around the allele. Since there are a lot of data points on the left side of the plot and only a few on the right side, we increased the radius of the circle from left to right from 0.1 to 0.5 linearly.

#### Analysis of conserved/non-mutable sites

120 complete mitochondrial genome sequences of mammalian species were downloaded from NCBI. The list of all species and accession numbers are provided in Table S2. Fasta files were aligned using MUSCLE online tool^15^. The alignments were parsed to find mtDNA sites that were identical between all 120 species.

### Modeling

To model accumulation of *de novo* somatic mtDNA mutations in hepatocytes we used an evolutionary simulation framework *SLiM*^16^ (v3.6). We regarded mtDNAs as individuals and all mtDNAs within a single cell as a population. Modelling was done following authors recommendations^17^ (section 14.9). The following parameters were used:

1. mtDNA reproduce clonally.
2. The recombination rate was set to zero.
3. Population size was kept constant and set to the number specified in a figure or figure legend. Generally, population size of 10,000 genomes was used for modeling mouse hepatocytes and population size of 5,000 genomes was used for modeling human hepatocytes.
4. The half-life for mtDNA in rat liver was estimated to be 9.4 days^18^ which translates into 78 generations over 2 years (maximum age of analyzed mice) and 3145 generations over 81 years (maximum age of analyzed human samples). We assume that in mouse and human the half-life of mtDNA is similar to what was measured in rat and, for convenience, round it to 80 and 3000 replacements for 2-year-old mouse and 81-year-old human, respectively.
5. Mutation emergence rate and selection coefficient varied in the models and are specified in figures and/or figure legends. Note, that this model does not include back-mutations. Hence, simulations using high mutation rates will be progressively less accurate.
6. To model the emergence, disappearance, and accumulation of a specific mutant allele, we simulated one site per genome and ran the simulation many times (Figures 2C, 3B, 3C, 4D, S4, S6C, S6D). To model NS/S across abundance intervals, the genome was limited to 11,338 sites which corresponds to number of protein coding sites in mouse mtDNA (Figures 6C and 6D). To model accumulation of mutations in the whole genome, genome size was set to 16,299bp (Figures 2D-2F). To simulate accumulation of a specific mutant allele at the tissue level we first simulated accumulation dynamics of the allele in 10,000 single cells and then computed the average abundance of the mutant allele across all simulated cells (Figures 4E and 4F).

#### Parameter space exploration

To find parameters that best describe the behavior of recurrent NCR mutations in 24-month-old mice we searched the parameter space. Specifically, we ran 315 models where mutation emergence rate varied from 10^-9^ to 10^-2^ and selection coefficient varied from −0.25 to +0.25.

Each simulation was run for 80 generations, population size was set to 10,000 genomes and it was repeated 9,833 times to match number of sequenced cells. For simulated data we have a record of all mutations present in a modeled cell, however in real sequencing data mutations present at levels below detection capability could not be detected. To mimic the observed data, each of 9,833 simulated cells was randomly assigned coverage of one of the cells from the experimental dataset. Then, simulated mutations present at levels below a sensitivity cutoff were set to zero. For each NCR mutation examined a sensitivity cutoff was calculated as 100% x 5/(assigned coverage at this site), where 5 is a minimum number of reads supporting the mutation. The resulting abundance distributions from each of 315 models were compared to the observed distribution of a mutation. First, the parameter sets that produced 2 time more or 2 times fewer positive cells than was observed were excluded. Next, we assessed the statistical significance of differences in abundance means between simulated and observed data using a permutation test. To calculate mean we used only cells/simulations that were positive for a mutation. 1,000 permutations were run to obtain a p-value. Abundance distributions produced with parameter sets close to the set with the highest p-value were inspected visually to identify parameters producing the best data-matching distributions.

#### Simulation of NS/S and STOP/S distribution for random emergence of mutations

To estimate NS/S and STOP/S expected for random mutagenesis of mtDNA, mutations in mouse (NC_005089) and human (NC_012920) mtDNAs were simulated using *Mutation-Simulator*^19^ (v 3.0.1). The transitions to transversions ratios (Titv) measured for the mouse dataset presented in Figure 1C (Titv = 4) and human (Titv = 5) dataset presented in Figure S6A were used as an input parameter for *Mutation-Simulator*. On average, 1.3 mutations were generated per simulation. The NS/S and STOP/S ratios were calculated for mutations generated in 360 simulations. This was repeated 1,000 times to obtain the ratios distribution.

Statistical analysis and data visualization were performed in Matlab (v. R2019a) and R (v. 4.1.3). Sample sizes, statistical tests and p-values are indicated in the text, figures and figure legends.

## Data availability

The sequencing data have been deposited in the NCBI Sequence Read Archive under PRJNA1146058.

---

**Figure S1.**
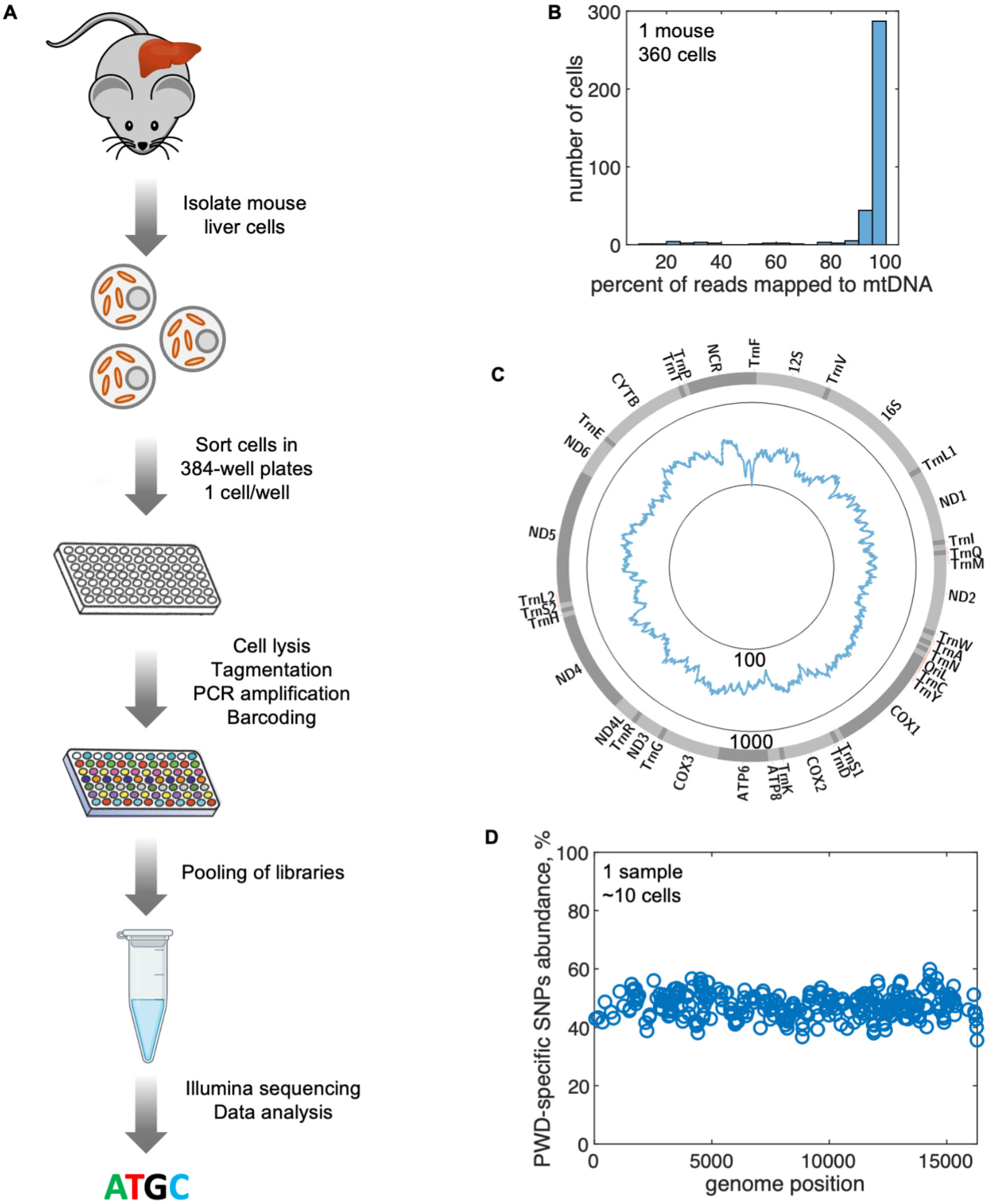
Plate-based single cell mtATAC method workflow and performance, related to Figure 1. (A) Schematic of the plate-based single cell mtDNA sequencing workflow. (B) Histogram showing the percent of raw reads mapping to mtDNA as measured for 360 cells isolated from 24-month-old C57BL6/J WT mouse liver. The majority of reads map to mtDNA. (C) Average mtDNA coverage as measured for 360 cells isolated from 24-months-old C57BL6/J WT mouse liver, log10 scale. While this method allows profiling the whole mtDNA, coverage is not uniform across the genome. Coverage in NCR is notably lower than in other regions of mtDNA. (D) Control showing the assessed abundance of PWD-specific SNPs in a sample where equal number of C57BL6/J and mtPWD hepatocytes were mixed (n =< 5 cells per sample type, the uncertain cell number is due to inefficient FACS sorting). This experiment shows the level of precision of mutation abundance detection.

**Figure S2.**
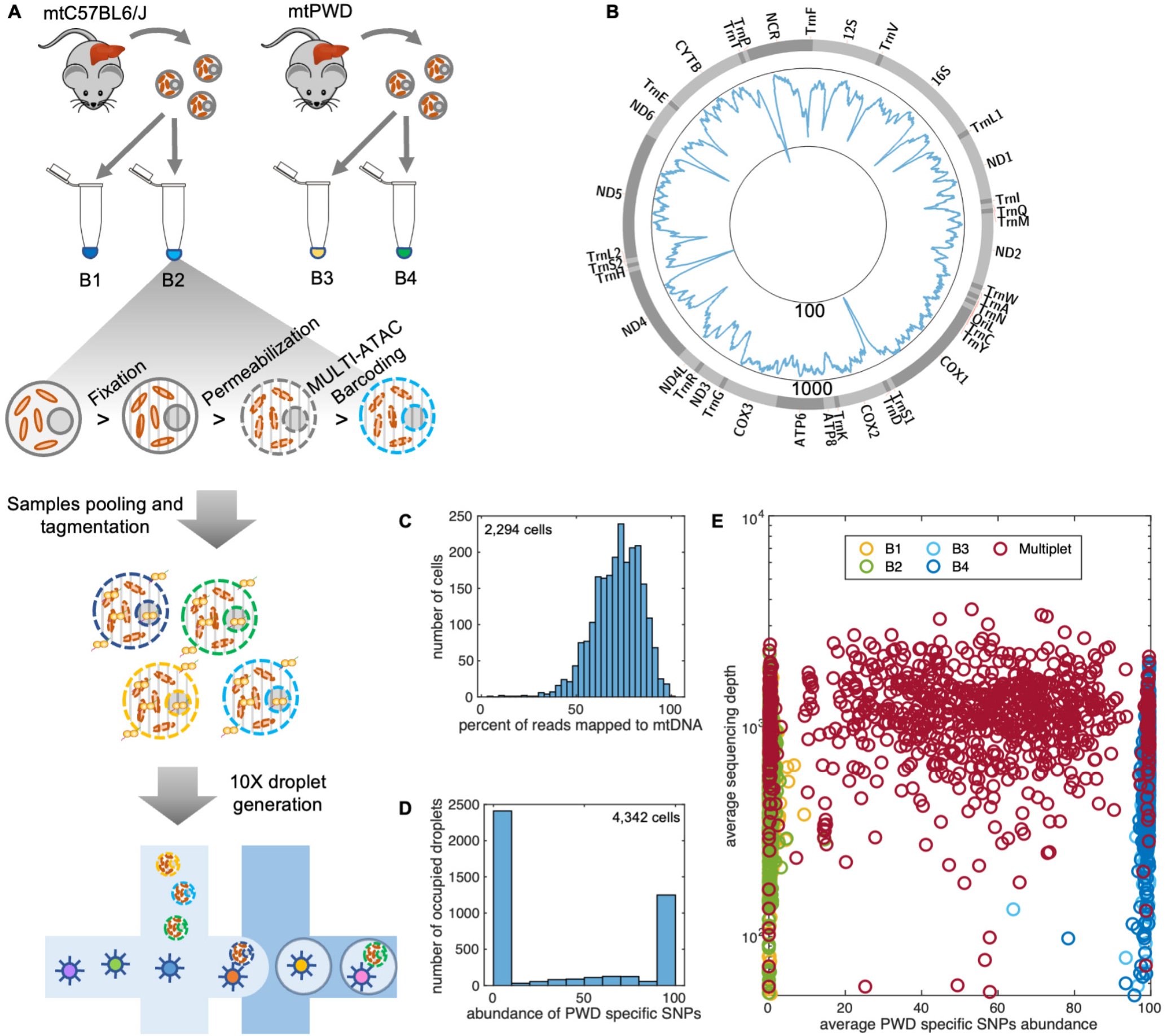
Single cell MULTl-1OX-mtATAC sequencing for profiling of somatic mtDNA mutations, related to Figure 1. (A) Schematic of the proof-of-principal experiment. In the experiment we mixed hepatocytes from two mouse lines with distinct mtDNA sequences (mtC57BL6/J and mtPWD). Samples were fixed in 1% PFA and permeabilized. Each biological sample was split in two halves and each of the four resulting samples was labeled with a unique lipid-tagged oligo (MULTI-ATAC barcodes B1-4). Next, all samples were pooled together and tagmented. After tagmentation, the cell suspension was loaded on 10X chip to generate 10X-mtATAC and MULTI-ATAC barcode libraries. (B) Average mtDNA coverage (read depth) in single mtC57BL6/J cells, n= 2294, 10910 scale. (C) A histogram showing enrichment of mtDNA reads over total reads in single mtC57BL6/J cells. (D) A histogram of the number of occupied droplets carrying different percentages of PWD-specific SNPs when mtC57BL6/J and mtPWD liver cells were mixed. The peak on the left represent droplets carrying mtC57BL6/J cells, and the peak on the right represents droplets carrying mtPWD cells. A shallow and wide distribution in the middle represents droplets with more than one cell (multiplets) carrying both mtC57BL6/J and mtPDW cells. (E) Classification of droplets according to sample-specific MULTI-ATAC barcodes matches cells’ mtDNA genotype. Droplets marked only by B1 or B2 align at the left consistent with mtC57BL6/J cells, while droplets marked with B3 or B4 align at the right consistent with mtPWD cells. There are three kinds of multiplet droplets: homotypic ones carrying both B1 and B2, which have only mtC57BL6/J cells; homotypic ones carrying B3 and B4 which have only mtPWD cells; and heterotypic ones carrying either B1 or B2 together with either B3 or B4, which carry both types of cells. The heterotypic multiplets with mixed mtC57BL6/J and mtPWD genomes are distributed across the middle of the graph, while the homotypic multiplets align as expected on the left or the right. Homotypic multiplets on the sides of the graph are masking underlying single cell data. B1: n=1052, B2: n=1208, B3: n=465, B4: n=696, Multiplets: n=921. Mean PWD-specific SNPs abundance for mtC57BL6/J cells is 0.35%, mean PWD-specific SNPs abundance for mtPWD samples is 99.12%.

**Figure S3.**
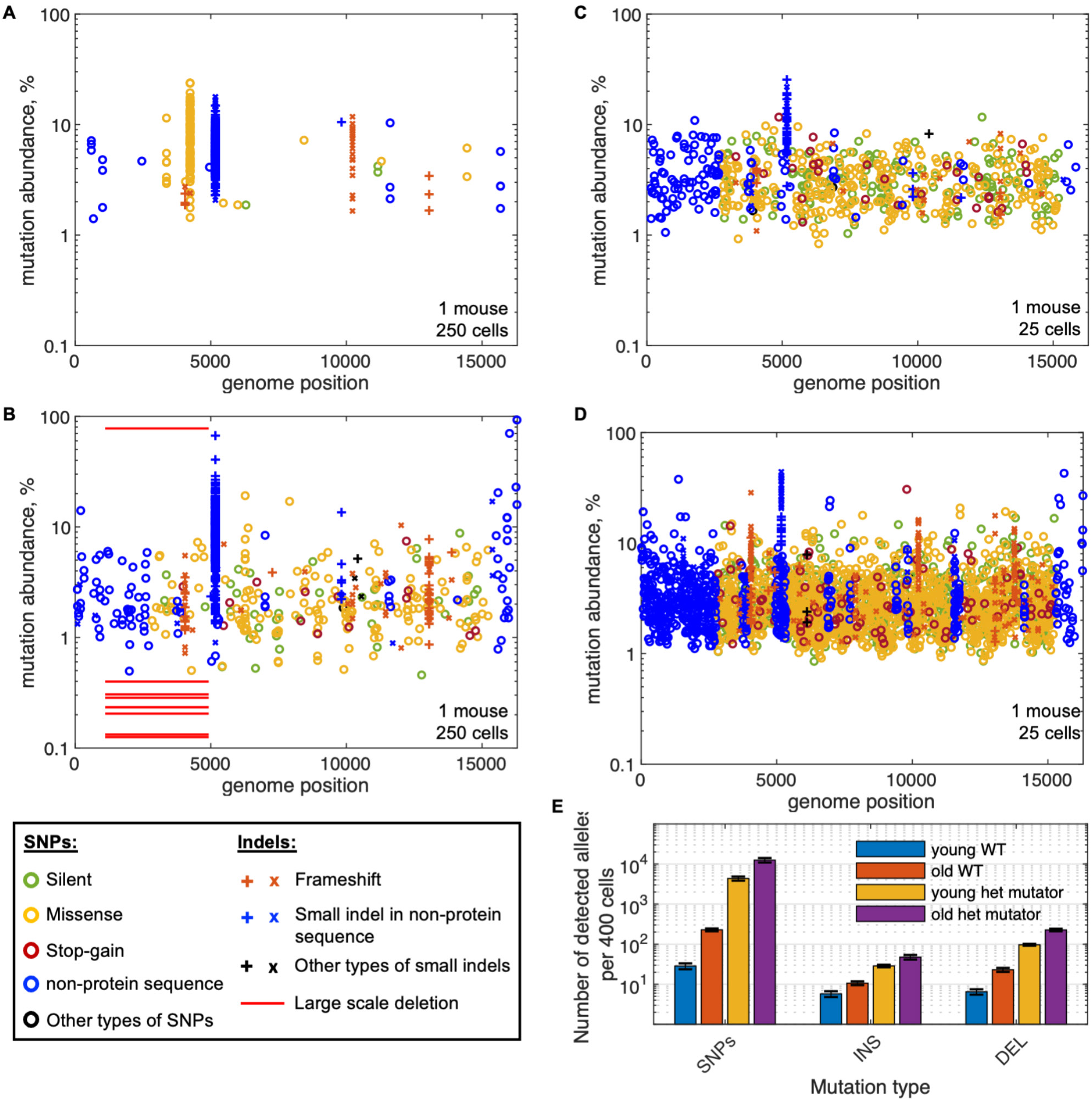
Spectrum of mtDNA mutations in *WT* and heterozygous mutator mouse livers, related to Figure 1. (A) Plot of all mutations detected in 250 cells isolated from liver of 3-months-old C57BL6/J WT mouse. Y-axis is logarithmic. Such representation allows better visualization of low abundance mutations. Note that the the overall absence of data towards the bottom of the distribution (below 1%) reflects the limitation in the sensitivity of detection of mutations and is not a true reflection of the frequency of mutations with lower abundances. (B) Plot of all mutations detected in 250 cells isolated from liver of 24-months-old C57BL6/J WT mouse. This is the same dataset that is presented in Figure 1C. Large-scale deletion in the minor arc was present in many cells at very low levels and in one cell at 80%. Other large-scale deletions detected in the presented example did not pass the threshold. (C) Plot of all mutations detected in 25 cells isolated from liver of 3-months-old heterozygous mutator mouse. (D) Plot of all mutations detected in 25 cells isolated from liver of 24-months-old heterozygous mutator mouse. (E) Number of detected mutant alleles per 400 cells of liver samples reported separately for SNPs, small insertions (INS) and small deletions (DEL). Samples from 4 young (3-months-old) and 3 old (24-months-old) C57BL6/J WT mice and 3 young and 3 old heterozygous mutator mice were analyzed. For this analysis data were subsampled to 100,000 chrM mapped reads per cell. This approach allows for fair comparison between samples but reduces the detection power of low abundance mutations. Fold change between number of detected alleles in WT and mutator mice presented in the main text were obtained from comparison of data from young animals. Number of detectable mutant alleles increased with age. Data in A-D were generated with plate-based approach. Data in E were generated with 10X-based approach.

**Figure S4.**
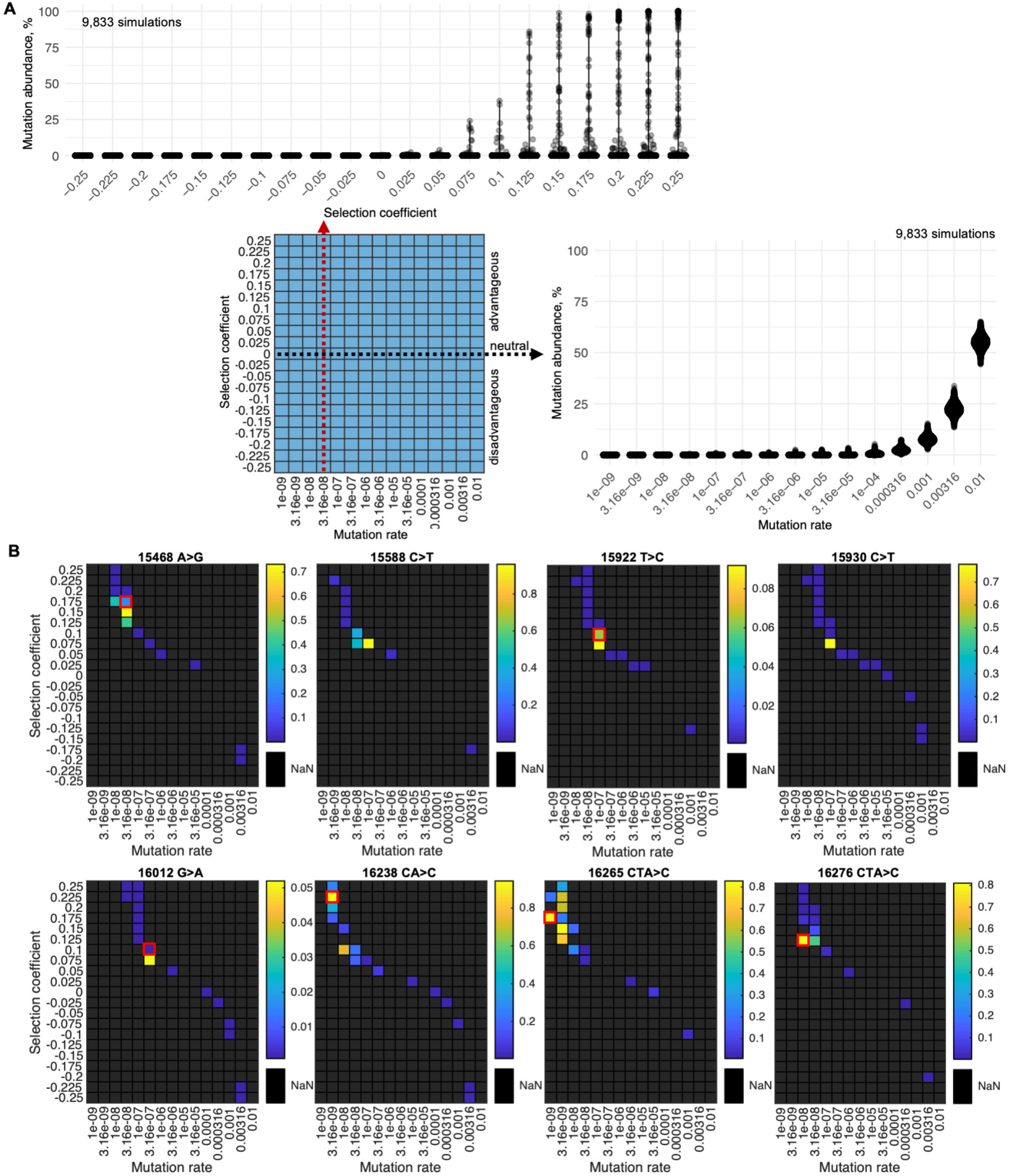
Modeling recurrent NCR mutations in WT mouse liver, related to Figure 4. (A) The impact of a range of mutation rates and selection coefficients on mutant allele abundance distribution among cells was explored in simulations. Top panel shows how selection coefficient impacts abundance distribution of a mutant allele emerging at 3.16×10-8per base pair per generation rate. Right panel shows how mutation rate impacts the abundance distribution of a neutral mutant allele. The results are shown for simulations that ran for 80 generations that we estimate to be equivalent to mtDNA turnover over 2 years of mouse life. Number of mtDNAs was set to 10,000. (B) Heatmaps showing similarity between abundance distribution of specified driver alleles and simulation outcomes. Colors represent “similarity value” (mean difference p-value based on 1,000 permutations) with yellow shades signifying the strongest similarity (see methods for details). Black squares represent parameter sets that produced 2-fold more or 2-fold less cells with detected mutant allele than real data. N=9833 cells/simulations. Abundance distributions produced with parameter sets close to the set with the highest “similarity value” were inspected visually. Red squares mark the model parameters that match best the data for the specified mutant allele based on visual inspection. Simulated mutant allele abundance distributions among cells produced with these parameters are presented in Figure 3D.

**Figure S5.**
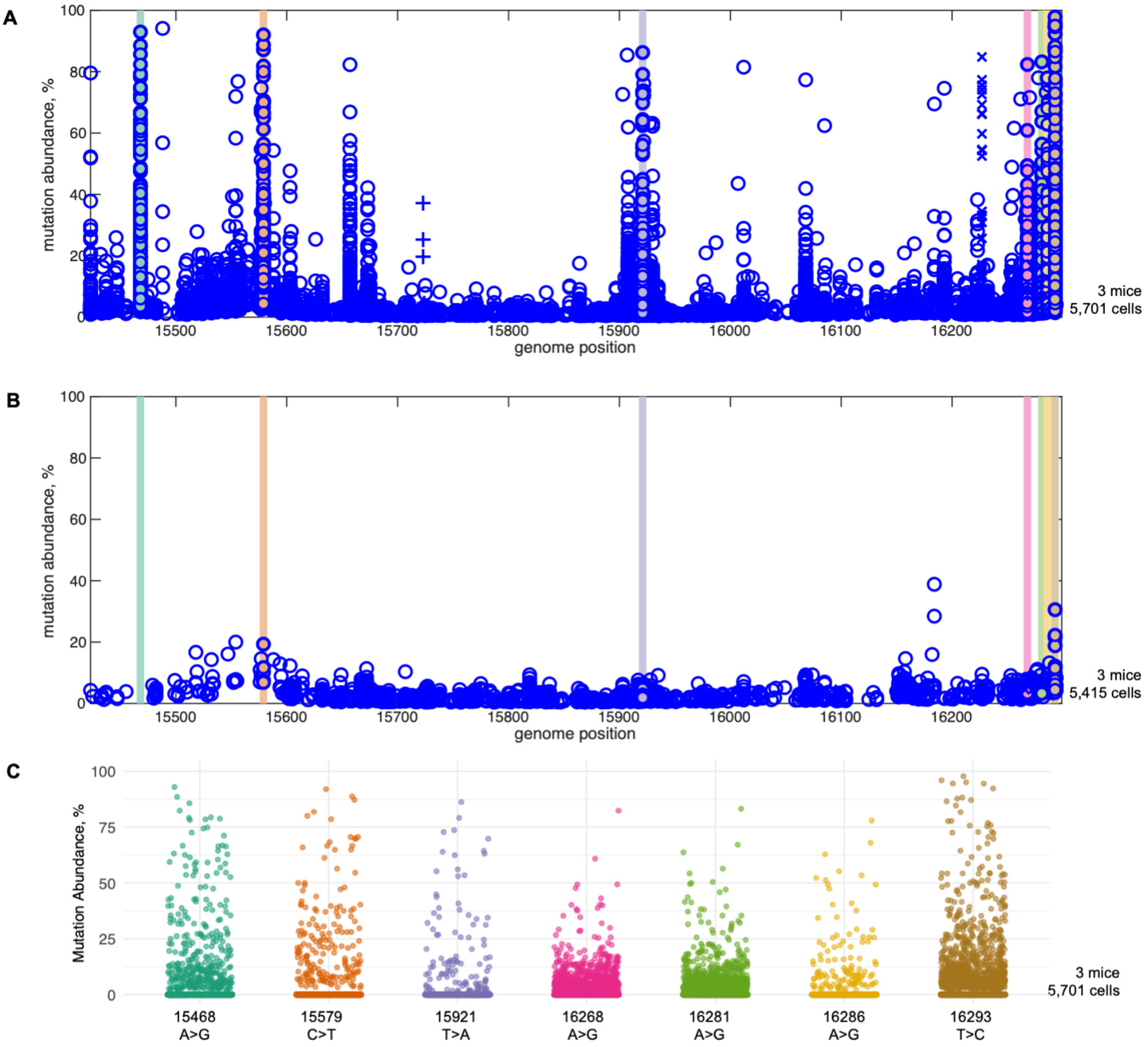
Specific mutations in NCR of heterozygous mutator mice confer a competitive advantage, related to Figure 4. (A) Spectrum of mutations in NCR of 24-months-old heterozygous mutator mouse liver cells. N=5,701 cells from 3 mice. Mutations that were present above 20% in at least 3 cells in all 3 mice are highlighted with color bars and marker filling. (B) Spectrum of mutations in NCR of 3-months-old heterozygous mutator mouse liver cells. N=5,415 cells from 3 mice. (C) Abundance distribution of the specified NCR mutations among liver cells of 24-months-old heterozygous mutator mice presented in panel **A.** Note a peculiar distribution of abundance of 16268A>G and 16281A>G mutations: these mutations are detected in many cells, however in majority of cells these variants stay below 10% and only in a few cells they climb to high levels. One possible explanation for such atypical abundance distribution is that these variants have selective advantage only in a small subset of liver cells.

**Figure S6.**
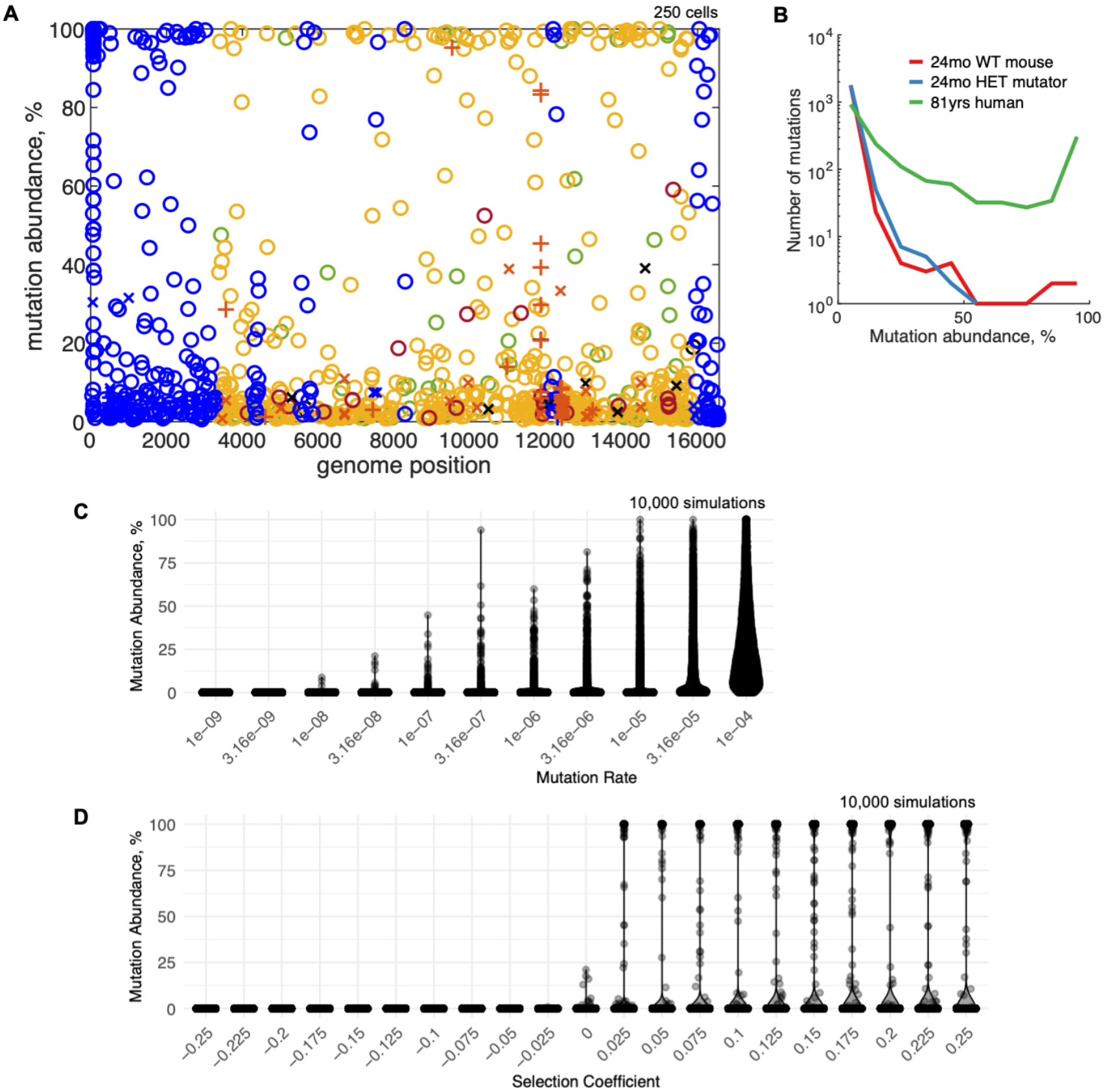
Spectrum of mtDNA mutations and modeling of mtDNA mutation accumulation in old human liver cells, related to Figure 7. (A) A spectrum of mtDNA mutations identified in 81-year-old human hepatocytes. (B) Abundance distribution of mutations in 24-month-old WT mice (3 mice form one experiment), 24-month-old heterozygous mutator mice (3 mice from one experiment) and 81-year-old human (one human) hepatocytes. Data from each sample were subsampled to equal number of reads per cell (100,000). For mouse data clones and mutations at p5171 (Oril) were excluded as they are very frequent and mask the signal from other mutations. Samples were normalized to have equal number of mutations: 1,829 mutations were randomly selected for each sample. These data indicate an increased mutation frequency has little influence on the proportion of mutations reaching high abundance, while age appears to have a large impact. (C) Impact of the mutation rate on the abundance distribution among cells of a neutral mutant allele. The results are shown for simulations that ran for 3,000 generations that we estimate to be equivalent to mtDNA turnover over −80 years of human life. Number of mtDNAs was set to 5,000. (D) The impact of the selection coefficient on abundance distribution among cells of a neutral mutant allele. Model parameters: mutaton rate 3.16×1a.a per base pair per generation, 3,000 generations, 5,000 mtDNAs per cell.

**Figure S7.**
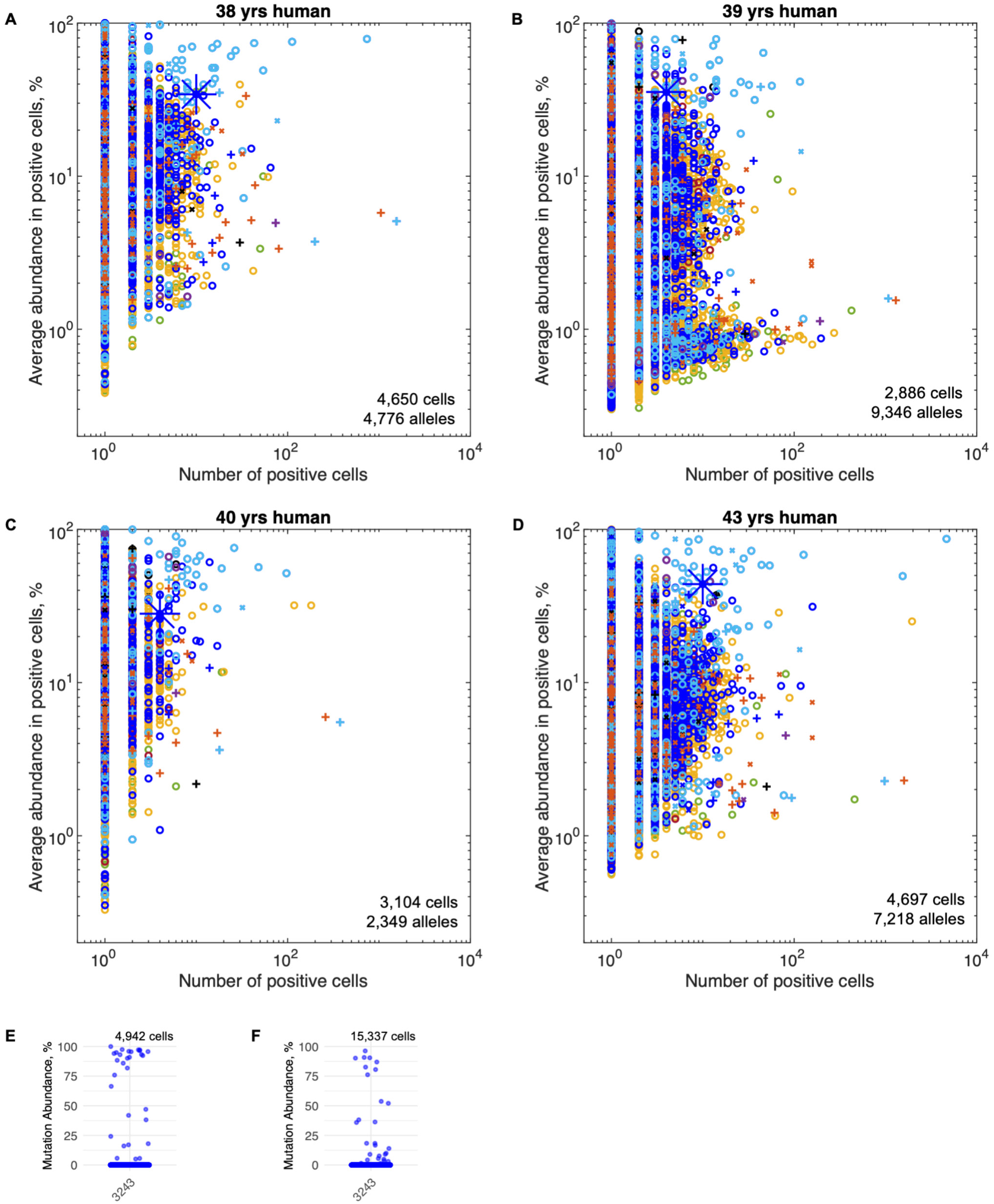
mtDNA mutation spectrums of middle-age human livers, related to Figure 7. (A-D) AAA vs #C plots for liver cells from middle-aged humans. 3243A>G allele colocalizes with NRC driver alleles and is marked by large blue*. The difference in number of alleles with low **AAA** among samples is due to difference in coverage. (E) Abundance distribution of 3243A>G mutation in liver cells of 41-year-old human presented in Figure 7. (F) Abundance distribution of 3243A>G mutation in liver cells of four near 40-year-old humans presented in (A-D).

